# Single-Cell Metagenomics Links Plasmid-Derived Antimicrobial Resistance to Hosts in the Pig Gut Microbiome

**DOI:** 10.64898/2026.07.24.740480

**Authors:** Alexander Zubov, Håkan Vigre, Saria Otani, Meilee Ling, Vibe Dalhoff Andersen, Marie Steengaard, Juliana Assis, Frank M. Aarestrup, Leonie Johanna Jahn, Alberto Santos, Patrick Munk

## Abstract

Antimicrobial exposure can alter gut resistance reservoirs, but bulk metagenomics alone often cannot distinguish whether observed changes reflect expansion of bacterial hosts, altered abundance of plasmid-derived sequences, or redistribution of mobile elements across host backgrounds. Here, we combined longitudinal bulk short-read metagenomics with selected bulk long-read and single-cell shotgun metagenomic sequencing to analyse faecal samples from six Danish pigs over 11 weeks, including an unplanned tiamulin exposure affecting the three pigs housed on the right side of the stable. We constructed a catalogue of 885 plasmid-derived sequences collapsed into 195 bins. Twenty-eight bins and 212 contigs carried resistance annotations, including ribosomal-target markers relevant to pleuromutilin exposure. Single-cell evidence linked subsets of plasmid-derived bins and contigs to bacterial host taxa, enabling host-resolved inspection of resistance-associated plasmid-derived features in longitudinal bulk metagenomes. The microbiome-wide plasmid-derived-sequence prevalence screen identified two bins with post-event associations, whereas resistance-gene abundance and host-attributed plasmid-derived-sequence abundance screens identified no significant host-resolved associations. Because exposure was unplanned and confounded with pen side and disease signs, treatment-response results are exploratory. The main contribution is a single-cell-informed microbial ecology workflow for linking plasmid-derived resistance features to host backgrounds and longitudinal abundance patterns in complex gut

## Introduction

Microbial communities are dynamic ecological systems in which community composition, encoded traits, and mobile genetic elements jointly shape ecosystem function in environmental, animal-associated, and human-associated habitats [1], [2]. Shotgun metagenomics has become central to microbiome research because it can profile taxonomic composition, functional potential, and reconstructed metagenome-assembled genomes without cultivation [3]. These genome-resolved approaches can link community members to metabolic or resistance-associated functions, but they remain limited when genes of interest occur on mobile elements whose host background cannot be inferred from bulk abundance alone [4].

Microbiomes also contain viruses and mobile genetic elements, including plasmids, which can move genetic information across bacterial lineages [5]. This is especially relevant for antimicrobial resistance (AMR) because resistance genes located on mobile elements can spread across host backgrounds, whereas chromosomal resistance genes mainly follow the abundance and transmission of their host lineage [5].

Standard metagenomic approaches that recover bulk genomes can fail to recover a substantial amount of plasmid diversity or other MGEs [6]. Long reads offer considerable advantages for recovering longer and often circular contigs that can be attributed to plasmids, and they can therefore complement short-read approaches in plasmidome reconstruction [7]. At the same time, plasmid recovery remains sensitive to sample preparation, input DNA, sequencing depth, abundance, and plasmid size; in particular, small or low-abundance elements can still be underrepresented [8]. Most importantly, standard bulk methods can study plasmids in the microbiome overall but cannot directly assign them to specific hosts. Proximity-ligation approaches such as Hi-C can link plasmids, antimicrobial resistance genes, and host chromosomes in complex communities, and have been used to connect the resistome and plasmidome to the microbiome [9]. However, these links are still inferred from physical proximity and are sensitive to sample structure, abundance, and linkage evidence, so they complement rather than replace cell-resolved approaches [10].

Single-cell sequencing is increasingly applied to microbiome research [11]. By separately barcoding and sequencing individual cells, sub-strain diversity can be uncovered that would otherwise be obscured by bulk methods [12]. Approaches for obtaining single-cell resolution commonly use droplet microfluidics [13] or semi-permeable capsules [14] to obtain single-amplified genomes (SAGs). Mining such genomic assemblies and recovering plasmids then allows analysis of the plasmidome with cellular context, as has recently been shown [15]. Although these studies showcased the potential of single-cell methods to resolve plasmid-host linkage, single-cell data can suffer from high dropout of genomic information, uneven and sometimes low coverage, fragmented assemblies, and taxonomic bias [16]. In essence, no single sequencing layer is capable of fully resolving the mobile resistome.

These limitations are especially relevant in complex environments such as the gut microbiome, where an increase in ARG abundance can reflect host expansion, plasmid abundance change, or redistribution of plasmids across bacterial hosts. A host-resolved analysis is therefore needed to better characterize the dynamics of the mobile resistome and, in particular, to identify candidate host backgrounds in which mobile ARG patterns occur.

We therefore combined longitudinal bulk short-read metagenomics with selected long-read and single-cell metagenomic sequencing of faecal samples from Danish pigs. Single-cell evidence was used to link plasmid-derived sequence (PDS) bins and contigs to bacterial host taxa, and these assignments were then transferred to longitudinal bulk metagenomes. The study is framed as a methods-oriented microbial ecology analysis with a natural tiamulin exposure test case, and treatment-response patterns are therefore interpreted as exploratory rather than causal.

## Materials and Methods

### Study Design

Faecal samples were collected from six pigs reared on a conventional Danish pig farm identified through the VETFORLIG research project. The stable section contained 24 pens arranged in two rows separated by a central access path, with approximately 25 pigs per pen. At the first visit, a few days after the pigs had entered the cleaned section at 10-12 weeks of age and approximately 30 kg body weight, three non-adjacent pens from each row were selected. One pig per selected pen was randomly chosen, ear-tagged, and spray-marked for repeated identification. Pigs B, D, and F were located on the left side of the central access path, whereas pigs A, C and E were located on the right side (figure 1A).

**Figure 1:**
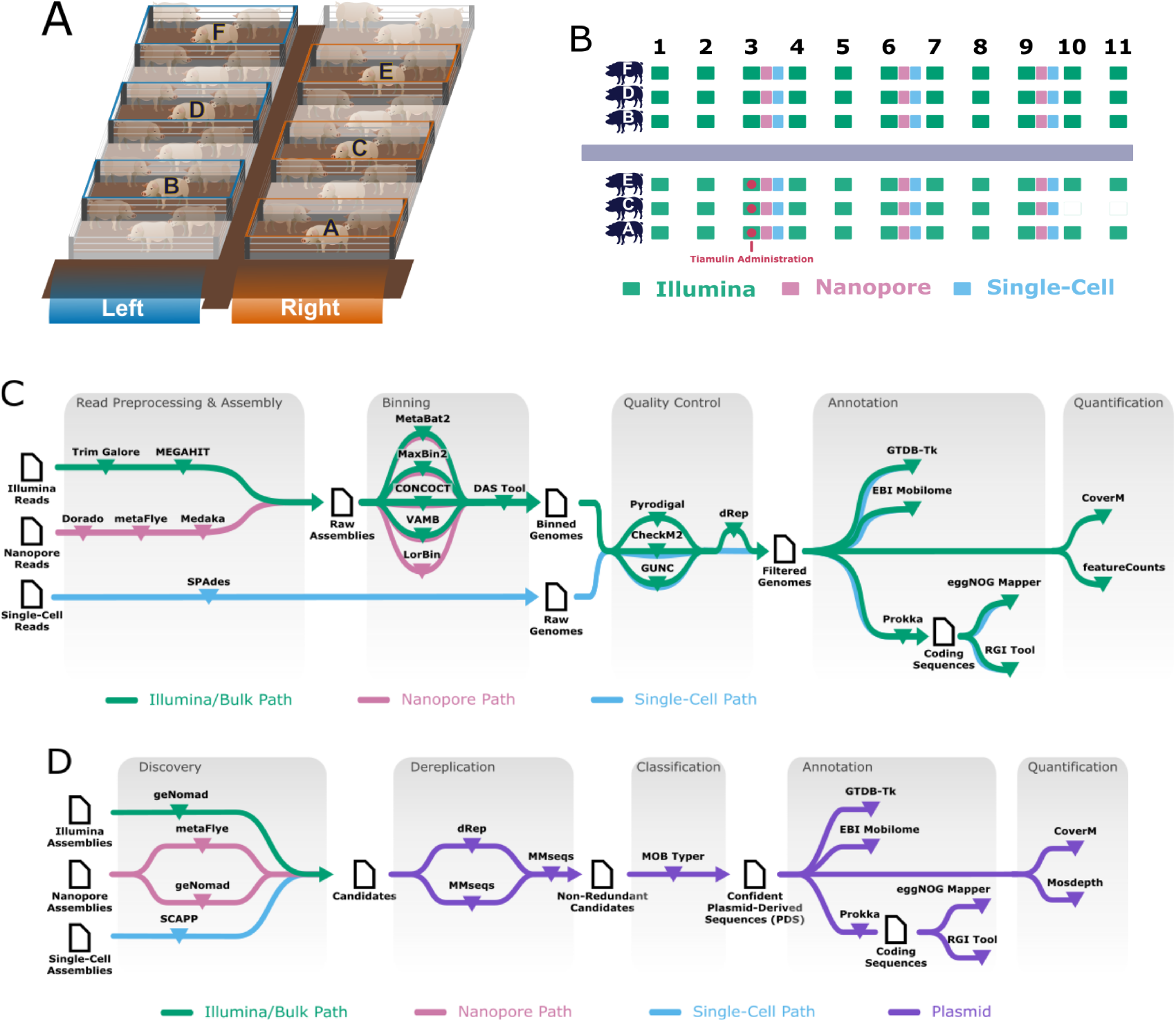
Study design and analytical workflow. (A) Schematic layout of the pig stable section. Six pigs were selected from non-adjacent pens, with pigs B, D, and F on the left side and pigs A, C, and E on the right side of the central access path. (B) Sampling schedule across the 11-week study. Green indicates bulk short-read shotgun metagenomic sequencing, magenta indicates bulk long-read shotgun metagenomic sequencing, and blue indicates single-cell short-read shotgun metagenomic sequencing. The tiamulin exposure event occurred between weeks 2 and 3 in the right-side pens. (C) Workflow used to construct the bulk and single-cell genome catalogues. (D) Workflow used to recover, filter, annotate, and quantify plasmid-derived sequences from the combined sequencing layers.

The six pigs were sampled weekly for 11 consecutive weeks until slaughter at approximately 24 weeks of age and 100 kg body weight. The exception was pig C which reached the required body weight at an earlier time point and was thus sent to slaughter after 9 weeks. Bulk short-read shotgun metagenomic sequencing was performed for all samples. In weeks 3, 6, and 9, bulk long-read shotgun metagenomic sequencing and semi-permeable capsule-based, single-cell short-read shotgun metagenomic sequencing were also performed (figure 1B).

The antimicrobial exposure was not planned before the start of the sampling. Between sampling weeks 2 and 3, clinical signs of diarrhoea were observed in pig E. Consequently, all pens in that row (which included pigs A, C, and E) were treated with tiamulin (pleuromutilin-type antibiotic) via drinking water for 5 days at a daily dose of 8 mg/kg body weight. The event was retained as a natural perturbation test case but cannot be considered a randomized intervention.

### Sample Storage, DNA Extraction, and Sequencing

Fresh faecal material was collected from each marked pig at each sampling visit and subsequently, samples were transported to the laboratory in styrofoam boxes with ice and split into aliquots immediately once reaching the lab later in the day (2-3 hours). Aliquotes were stored at -80 °C until processing. Aliquots used for bulk short-read sequencing, bulk long-read sequencing, and single-cell sequencing were processed from different frozen aliquots.

For bulk short-read sequencing, DNA was extracted from 0.1 g of faecal material and after preparation, libraries were sequenced on Illumina NovaSeq 6000 to generate paired-end reads of 150 bp. For bulk long-read sequencing, high-molecular-weight DNA was extracted from 0.1 g of faecal material. Library preparation followed Ivanova et al. [17] with the following modifications: sequencing was performed on MinION platform using R9.4.1 flow cells and the ligation sequencing kit SQK-LSK109.

For single-cell sequencing, 0.1 g of each faecal sample was processed using the semi-permeable capsule (SPC)-based single-cell metagenomic workflow described by Ling et al. [14], with the following study-specific modifications. Cell detachment and encapsulation were performed using a customized single-cell microbial WGS kit (catalogue no. CKP-BARK8; Atrandi Biosciences). Approximately 10,000 cells from each sample were loaded at a target occupancy of lambda = 0.1 cells per SPC, corresponding to approximately 100,000 SPCs per sample. Under this design, most SPCs were expected to contain no intact cell. SPCs from multiple samples were pooled in each combinatorial barcoding run. Libraries were sequenced on an Illumina NovaSeq X Plus instrument to generate 150 bp paired-end reads. Barcode demultiplexing, read count filtering, assembly, and SAG quality filtering are described in Supplementary Methods.

The sample and run accessions as well as additional metadata are reported in Supplementary Table S1.

### MAG and SAG Catalogue Construction

The overall workflow is summarized in figure 1, with extended parameters provided in Supplementary Methods. Software versions and database releases used in the analysis are reported in Supplementary Table S2.

### Read Preprocessing and Metagenomic Assembly

Short-read Illumina data were quality-trimmed with TrimGalore and assembled with MEGAHIT [18]. ONT reads were converted from Fast5 to Pod5, basecalled and demultiplexed with Dorado using the dna_r9.4.1_e8_sup@v3.6 model, assembled with metaFlye, and polished with Medaka [19], [20], [21]. Single-cell reads were assigned to SPCs using the four-component combinatorial barcode scheme and were quality-trimmed and retained as described by Ling et al. [14]. Retained single-cell reads were subsequently assembled as in Kawano-Sugaya et al. [15] using SPAdes in single-cell mode with –careful enabled and repeat resolution disabled [22]. Bulk assemblies were generated at single-sample, within-pig, and whole-study levels, while short- and long-read assemblies were kept as separate evidence layers.

### Prokaryotic Genome Recovery, Dereplication, Annotation, and Quantification

Bulk contigs were binned with MetaBAT2, MaxBin2, CONCOCT, VAMB, and, for long-read assemblies, LorBin [23], [24], [25], [26], [27]. Bin candidates were refined with DAS Tool [28]. Genomes were assessed with CheckM2 and GUNC [29], [30]. MAGs were retained at >=75% completeness, >=80% coding density, and <=10% contamination. SAGs were retained at >=20% completeness, <=30% contamination, and <0.5 clade separation score. MAGs were dereplicated with dRep, whereas SAGs were not dereplicated [31]. MAGs and SAGs were taxonomically annotated with GTDB-Tk, and coding sequences were predicted with Prokka [32], [33]. Protein-level gene clusters were generated with MMseqs2 and annotated with eggNOG-mapper and RGI [34], [35], [36], [37], [38]. Bulk metagenomes and gene features were quantified with CoverM and featureCounts [39], [40].

### Single-Cell Processing

Single-cell data were processed in Python with SnapATAC2 for visual inspection and downstream feature analyses [41]. In this manuscript, barcode refers to the raw single-cell assignment layer, whereas cell refers to barcodes passing quality control and, where relevant, receiving a taxonomic assignment. Barcode, cell, and feature filtering are described in Supplementary Methods. Functional count matrices were embedded using Laplacian eigenmaps, clustered with Leiden clustering, and visualized with UMAP [42], [43].

### PDS Recovery, Annotation, Quantification, and Host Linkage

We use plasmid-derived sequence (PDS) for recovered sequences with evidence of plasmid origin without assuming that the recovered sequence represents a complete circular plasmid. Retained PDS contigs are sequence-level catalogue entries with the prefix PLS_. PDS bins are Pling-derived collapsed groups of related retained PDS contigs with the prefix PLB_. The two levels were used for complementary purposes. PDS contigs preserve sequence provenance, ARG localization, and fine-scale host-resolved inspection, whereas PDS bins reduce fragmentation of related sequences and provide a more conservative collapsed catalogue unit for catalogue summaries, co-occurrence analyses, and host-linkage summaries. This distinction is important because most recovered plasmid-derived sequences are expected to be incomplete and often originate from the same plasmid.

### PDS Discovery and Binning

Candidate PDSs were recovered from geNomad predictions on Illumina and ONT assemblies, circular metaFlye ONT contigs, and SCAPP predictions from single-cell assemblies [44], [45].Candidates were dereplicated, clustered with MMseqs2, and classified with MOB-Typer based on similarity to known plasmids and plasmid mobility markers [46]. Sequences at least 1,000 bp long and assigned to middle, high, or very high confidence were retained as PDS contigs. Retained PDS contigs were grouped into PDS bins with Pling [47]. Extended filtering, confidence classification, Pling, and unique-region parameters are provided in Supplementary Methods.

### PDS Annotation and Quantification

PDS contigs and PDS bins were annotated separately. PDS-contig annotation used Prokka-predicted coding sequences from retained PDS contigs followed by eggNOG-mapper and RGI. PDS-bin annotation used bin-level sequence outputs, with translated coding sequences redundancy-reduced with CD-HIT before eggNOG-mapper and RGI annotation [36], [37], [38], [48]. Bin-level annotations were used for collapsed catalogue summaries and ARG co-occurrence analyses, whereas contig-level annotations were used for ARG localization and host-resolved inspection.

Bulk short-read data were used to estimate PDS abundance at PDS-bin and PDS-contig levels. Reads were aligned to a combined PDS/MAG reference, and PDS depth was calculated over informative regions after masking sequence shared across PDS features and sequence shared with MAGs. Genome-normalized abundance was calculated as PDS depth divided by host species marker depth. Because PDS recovery is incomplete and coverage is estimated over informative recovered regions, this metric was interpreted as genome-normalized PDS abundance rather than literal complete-plasmid copy number.

### Single-Cell PDS Evidence

For single-cell PDS evidence, barcode reads were aligned to the retained PDS catalogue with BWA-MEM2 and quantified with Mosdepth [49], [50]. PDS-unique regions were defined using MUMmer/NUCmer self-alignments [51]. Regions shared with MAGs or the barcode’s own SAG assembly were subtracted before barcode-level quantification. PDS evidence in a barcode was classified as strong if breadth at 1x was >=0.30 or breadth at 3x was >=0.20, and weak if breadth at 1x was >=0.20 but did not meet the strong-evidence threshold.

### Host-Linkage Layers

Host linkage was inferred separately for PDS bins and PDS contigs. Barcode-level PDS evidence was joined to taxonomically assigned cells, and PDS-feature/species enrichment was tested using the SciPy implementation of a one-sided Fisher exact test followed by Benjamini-Hochberg correction [52], [53]. Stringent host assignments required at least two supporting cells and q < 0.05. All observed host-feature pairs were retained as a permissive layer. Stringent linkage was used for conservative host-linkage claims and visual summaries, whereas permissive linkage was retained for sensitivity analyses and exploratory abundance modelling.

## Results

### A Multi-Technology Approach Recovers a Diverse Pig Gut PDS Catalogue

We constructed a catalogue of plasmid-derived sequences (PDSs) from short-read, long-read, and single-cell assemblies. The workflow generated 1,181 dereplicated metagenome-assembled genomes (MAGs) and retained 6,533 single-amplified genomes (SAGs) after quality control. The PDS workflow retained 885 PDS contigs and collapsed related contigs into 195 Pling-derived PDS bins. Contigs preserved sequence provenance and antimicrobial resistance gene (ARG) localization, whereas bins reduced fragmentation and supported conservative catalogue-level summaries. Of the 195 bins, 128 were singletons and 67 contained multiple PDS contigs. Twenty-eight bins carried ARG annotations, and 11 had stringent single-cell host-attribution evidence (Supplementary Table S5, figure 2).

**Figure 2:**
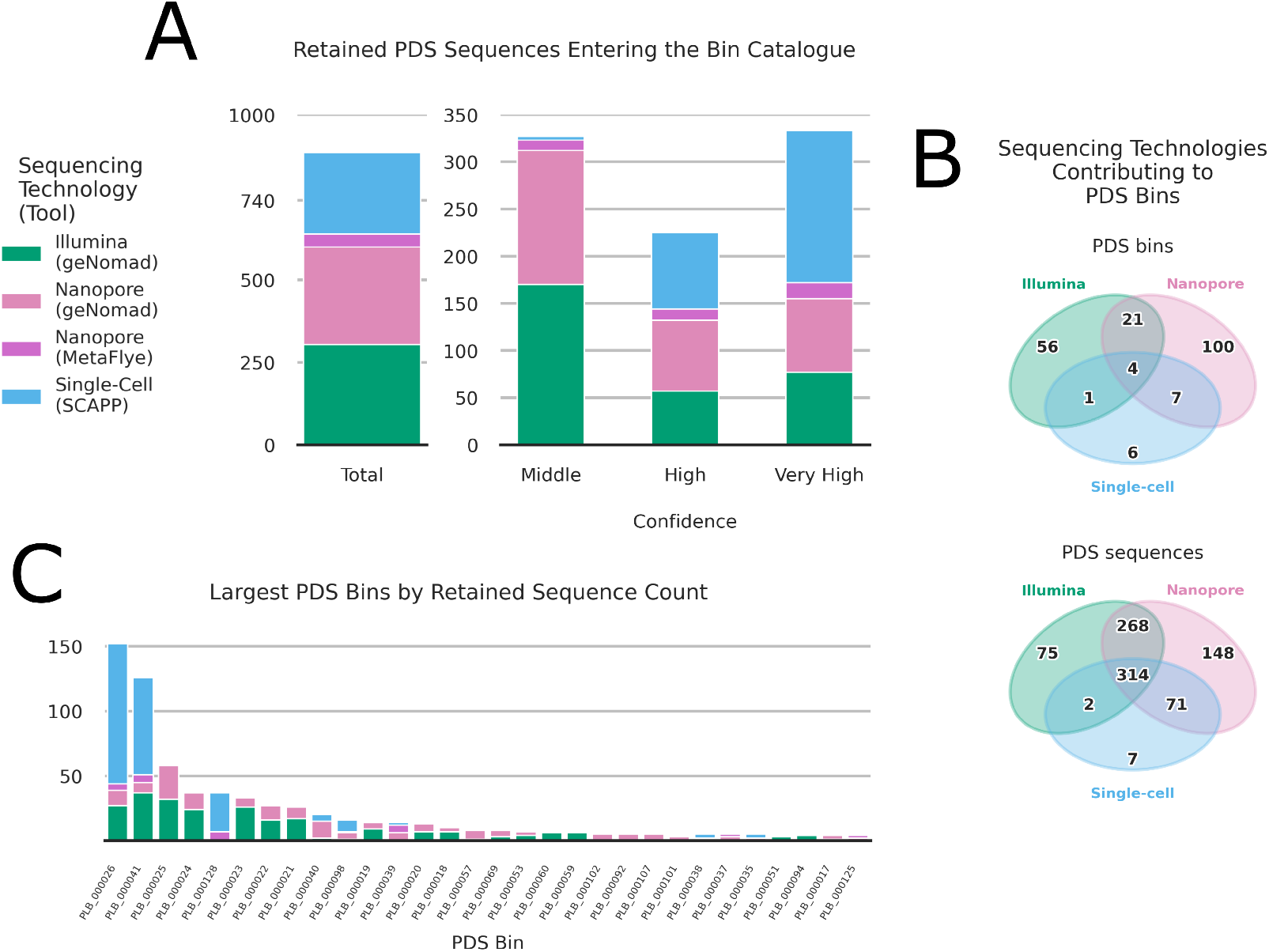
Multi-technology recovery of plasmid-derived sequences. (A) Contribution of sequencing technologies and prediction tools to the retained PDS-sequence catalogue, stratified by confidence class. Confidence classes were assigned based on similarity to known plasmids and the presence of plasmid marker genes. (B) Sequencing-technology support patterns for PDS bins and retained PDS contigs. (C) The 30 largest PDS bins, ranked by retained PDS-sequence count, with bars coloured by contributing sequencing technology and prediction tool.

The sequencing layers contributed complementary, only partially overlapping PDS evidence. Most retained PDS contigs originated from long-read or short-read geNomad predictions, whereas single-cell SCAPP and circular long-read metaFlye contigs contributed smaller fractions (figure 2A). Single-cell-derived sequences however formed the largest contribution to the very-high-confidence class. At the bin level, 33 PDS bins were supported by more than one sequencing technology and four by all three technologies (figure 2B). MOB-Typer classified 116 PDS bins as non-mobilizable, 77 as mobilizable, and two as conjugative [46] (Supplementary Table S5).

### ARG-Bearing PDSs Include Pleuromutilin-Relevant Resistance Markers

To understand the plasmid-derived resistome rather than the total community resistome, we screened PDS coding sequences for AMR annotations. RGI annotation identified 28 ARG-bearing PDS bins and 212 ARG-bearing PDS contigs. At the bin level, the most frequent ARG labels were *tet(W)* in 8 bins, *APH(3’)-IIIa* in 7, *ermB* in 6, and both *SAT-4* and *tet(40)* in 5, and *cfrE* in 3. The main ARG-count summary is shown at contig level to support localization and host-resolved inspection, whereas bin-level ARG counts are provided in Supplementary Figure S1 and Supplementary Table S4 (figure 3A; Supplementary Figure S1).

**Figure 3:**
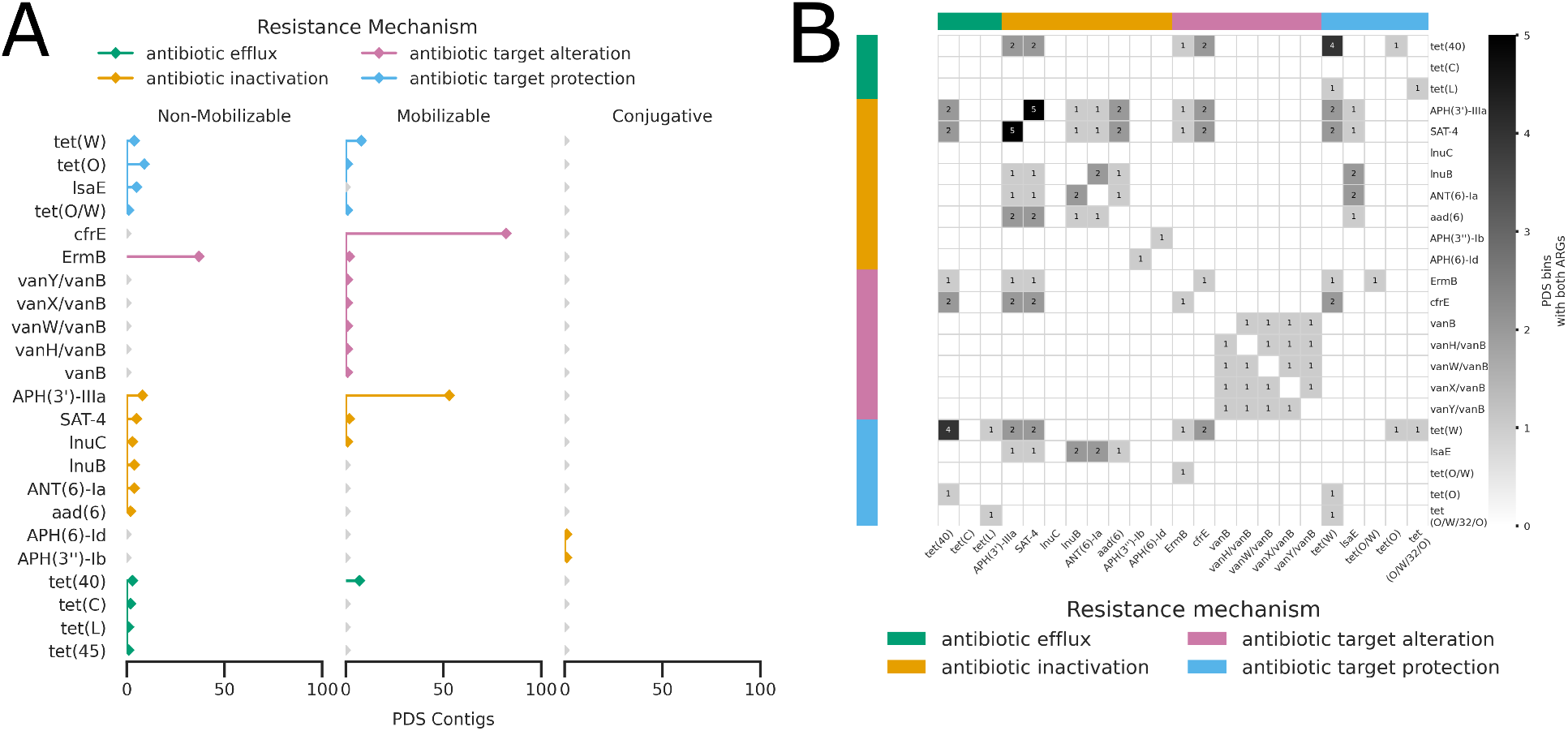
Antimicrobial resistance gene (ARG) content of the PDS catalogue. (A) Number of ARG-bearing PDS contigs carrying each ARG annotation, stratified by predicted PDS-contig mobility class and coloured by resistance mechanism. Bin-level ARG counts are provided in Supplementary Figure S1. (B) Co-occurrence of ARG annotations on the same PDS bin. Darker cells indicate more frequent co-occurrence and blank cells indicate no observed co-occurrence.

Because tiamulin was administered during the sampling period and acts at the peptidyl-transferase centre of the 50S ribosomal subunit, we specifically examined PDS-associated resistance markers affecting this ribosomal target [54], [55], [56], [57]. Most distinct ARG annotations represented antibiotic target alteration or antibiotic inactivation. Among the detected markers, *cfrE* had the clearest direct relationship to pleuromutilin resistance because Cfr-family methyltransferases modify 23S rRNA within the peptidyl-transferase centre [54], [55], [56]. *ermB* also modifies domain V of the

23S rRNA but is more appropriately interpreted as a broader antibiotic resistance determinant than as a tiamulin-specific marker 57. All three *cfrE* -bearing PDS bins were predicted as mobilizable, although *cfrE* represented only a part of the complete PDS-associated ARG catalogue.

In several cases, ARGs co-occurred with other ARGs. The strongest co-occurrence was *APH(3’)-IIIa* with *SAT-4*, found in 5 PDS bins (Figure 3B). Other co-occurrences included *tet(40)* with *tet(W)* in four PDS bins and *tet(40)* with *tet(O)* in one PDS bin. Pairs found in two PDS bins each included *lnuB* with *lsaE, cfrE* with *tet(W), SAT-4* with *cfrE*, and *APH(3’)-IIIa* with *cfrE. APH(3)-Ib* and *APH(6)-Id* occurred together in the same PDS bin predicted as conjugative. These genes correspond to the linked *strA–strB* streptomycin-resistance module originally described in the broad-host-range plasmid RSF1010 [58]. These patterns demonstrate co-carriage within collapsed PDS bins but do not establish that the linked genes were co-selected during the sampling period.

### Single-Cell Evidence Links ARG-Bearing PDS to Bacterial Species

We used PDS-unique regions to connect PDS bins and PDS contigs to individual single-cell barcodes. At the PDS-bin level, strong single-cell evidence comprised 9,584 observations across 4,216 cells and 92 PDS bins, including 10 bearing ARGs. Weak evidence comprised 2,151 observations across 1,619 cells and 64 PDS bins, including 9 ARG-bearing PDS bins (Supplementary Table S5). The PDS-contig layer was used as the primary ARG-localization layer for host-resolved inspection because contigs preserve finer sequence-level information than collapsed bins, whereas the bin layer provides a more conservative, collapsed view of related PDS evidence.

Most retained PDS detections were supported across substantial portions of the corresponding sequence, but many individual PDS features occurred in only a small number of cells. We therefore tested whether each PDS feature was enriched in particular bacterial species relative to its overall frequency across taxonomically assigned cells. This produced a stringent host-assignment set for conservative interpretation and a permissive set for sensitivity analyses. ARG-bearing features detected in the single-cell layer included 11 ARG-bearing PDS bins and 43 ARG-bearing PDS contigs after applying the current PDS–ARG annotations used for figure 4 and Supplementary Figure S2.

**Figure 4:**
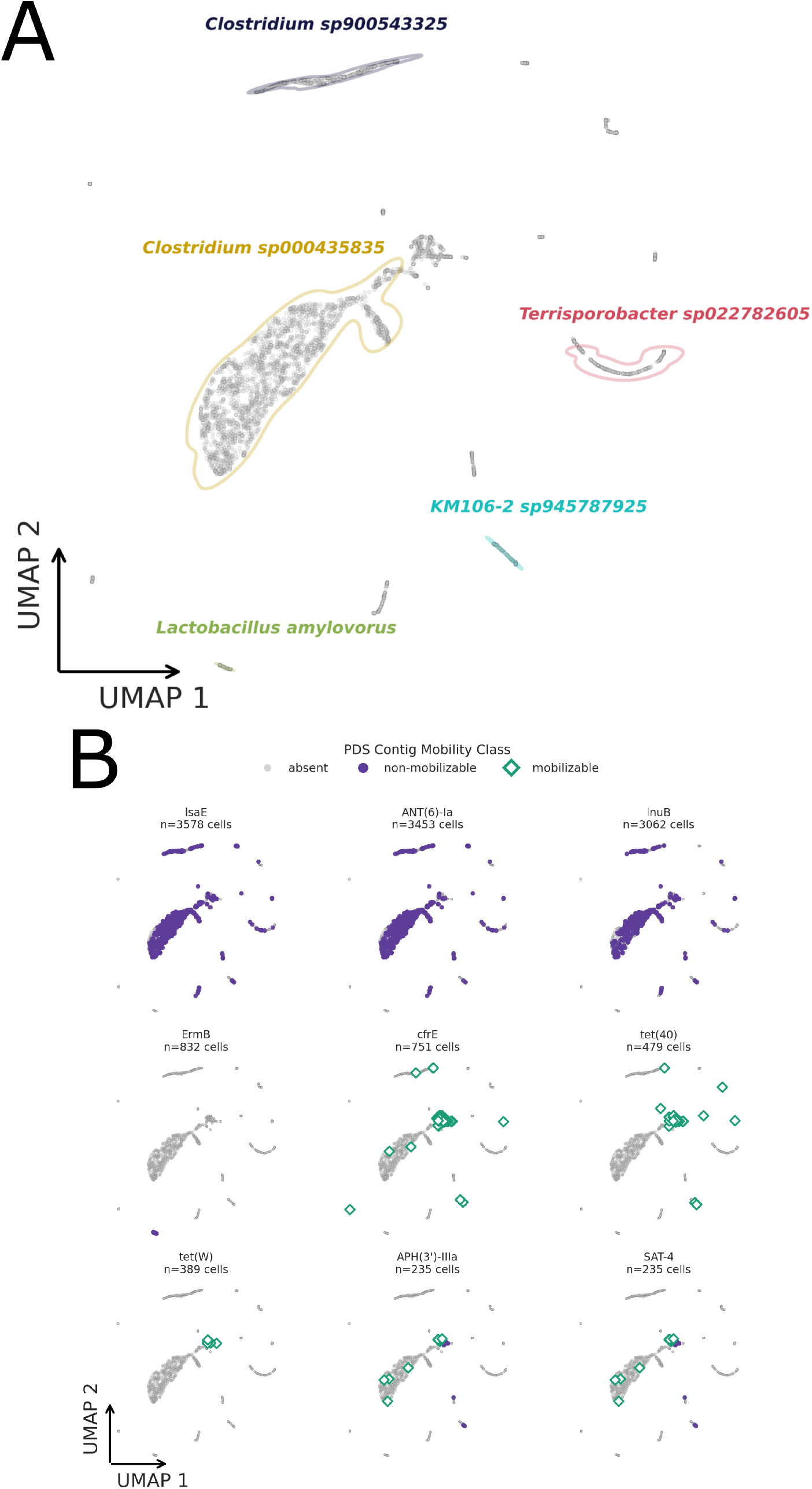
Single-cell evidence for host-associated ARG-bearing PDS contigs. (A) Uniform manifold approximation and projection (UMAP) of functional embeddings derived from the filtered single-cell genome catalogue. The embedding is based on functional annotations after filtering cells and gene features. The five species with the highest cell counts are highlighted. (B) Per-cell evidence for ARG-bearing PDS contigs, separated by ARG annotation and predicted PDS-contig mobility class. Grey points indicate cells without evidence for the displayed feature, filled points indicate evidence for non-mobilizable PDS contigs, and open diamonds indicate evidence for mobilizable PDS contigs. The value of *n* above each panel indicates the number of cells containing at least one PDS contig with the corresponding ARG annotation.

Functional embedding separated several of the most frequently recovered taxa, while ARG-bearing PDS-contig evidence occurred in restricted subsets of cells rather than throughout the embedding (figure 4). A corresponding bin-level view is provided in Supplementary Figure S2.

### Host Attribution Resolves a Subset of the PDS-Associated Resistome

Single-cell evidence connected multiple gut bacterial taxa to ARG-bearing PDS features. Stringent PDS-bin linkage yielded 17 host-bin pairs involving 11 bins and 11 host species, including three ARG-bearing bins. Stringent PDS-contig linkage yielded 67 host-contig pairs involving 36 contigs and 25 host species, including 13 ARG-bearing contigs. The contig-level analysis therefore preserved more ARG-specific host associations, whereas the bin-level analysis provided a more conservative collapsed view (figure 5, Supplementary Figure S3, Supplementary Tables S6 and S7).

**Figure 5:**
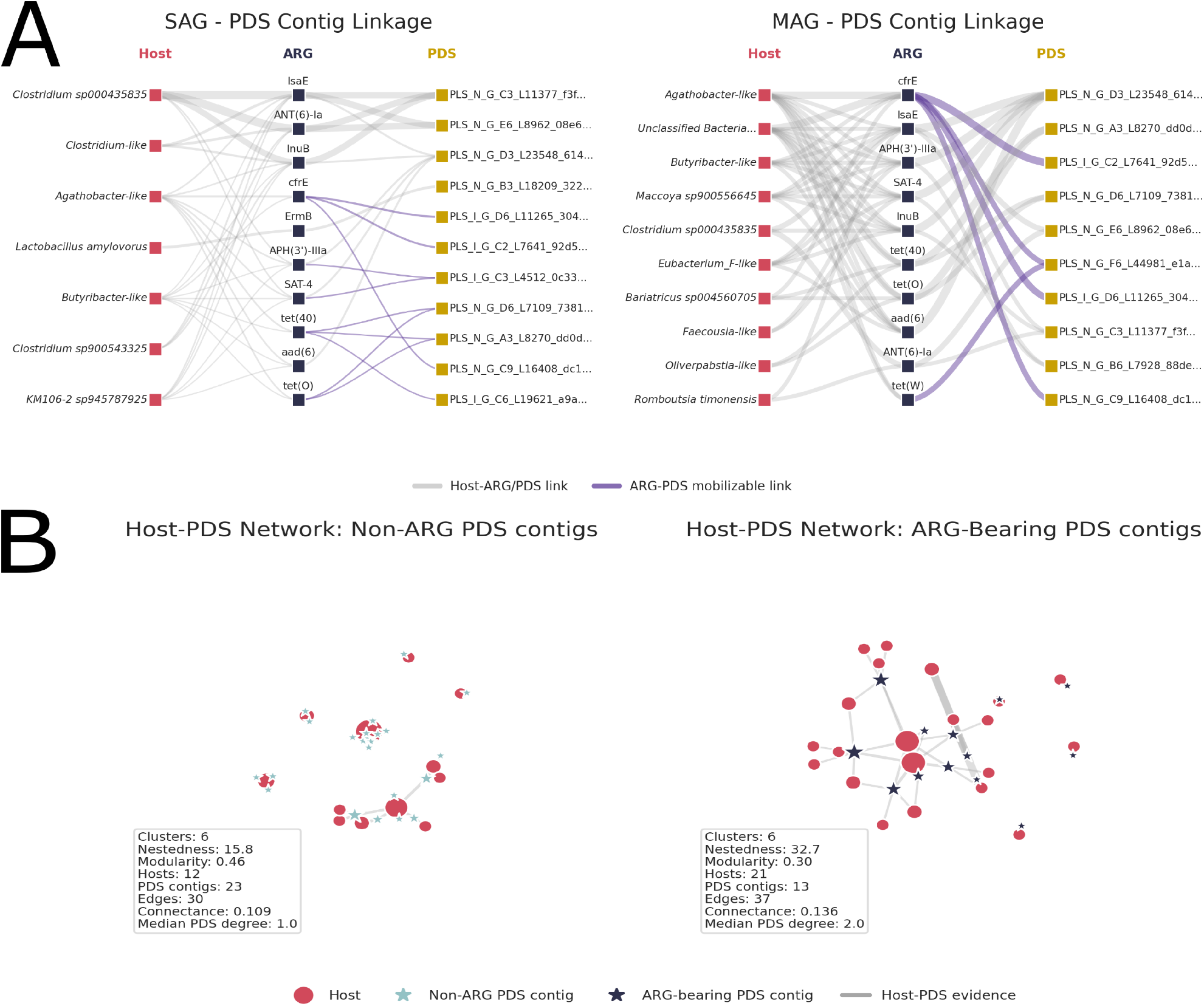
Host attribution of ARG-bearing PDS contigs. (A) Sankey-style diagrams showing associations between bacterial hosts, antimicrobial resistance genes, and PDS contigs. Link thickness summarizes the number of unique host-ARG or ARG-PDS attributions. Purple ARG-PDS links indicate that the PDS contig belongs to a PDS bin predicted as mobilizable. SAG-side plots (left) use strong single-cell evidence for displayed host-ARG-PDS triads, whereas MAG-side plots (right) use stringent host-support. (B) Bipartite host-PDS networks comparing non-ARG-bearing and ARG-bearing PDS contigs. Network edges were derived from host-linkage layers and are shown as exploratory descriptors of host-linkage graph structure.

ARG-bearing and non-ARG-bearing host–PDS networks differed descriptively in their observed connectivity (as observed by Risely et al. [59]), although these graph summaries were not subjected to inferential testing (figure 5B; Supplementary Figure S3C,D).

### Exploratory Models Identify PDS-Bin Prevalence Associations but No Host-Resolved Abundance Shift

Between weeks 2 and 3, the three pigs housed on the right side of the stable received tiamulin following the appearance of diarrhoea in pig E. We therefore tested whether PDS-bin prevalence or abundance changed after this event and whether the change differed between the right- and left-side pigs. Two PDS bins showed significant prevalence associations with the post-event period and/or the post-event-by-pen-side interaction. In contrast, no PDS-bin abundance, PDS-associated ARG-family abundance, or host-attributed PDS-contig abundance feature met the prespecified threshold of q < 0.05 and beta coefficient of 0.5. Broad MAG and order-attributed ARG screens and selected host-PDS trends are provided as contextual analyses (Supplementary Figures S4-9, Supplementary Tables S8 and S9). Because tiamulin exposure was unplanned and confounded with pen side and disease signs, all associations are interpreted as exploratory rather than as causal treatment effects.

## Discussion

This study presents a host-resolved workflow for analysing plasmid-derived resistance reservoirs in the pig gut microbiome. Longitudinal short-read metagenomics provided quantitative resolution, long-read sequencing improved the recovery of PDS contigs, and single-cell sequencing supplied host-linkage evidence. Together, these layers connected subsets of PDS bins and contigs to bacterial taxa and enabled their inspection across longitudinal bulk metagenomes. The principal contribution is therefore methodological and ecological: the workflow identifies candidate host backgrounds for plasmid-derived resistance features that cannot be assigned from bulk abundance alone. The unplanned tiamulin exposure provided an exploratory perturbation but did not support causal host-specific response inference.

Previous pig studies have shown that microbiome and resistome responses to antimicrobial exposure vary across cohorts, farms, drug classes, and experimental designs [60], [61], [62]. Such variation is expected because an increase in ARG abundance can result from expansion of the bacterial host, a change in mobile-element copy number within the host, redistribution of an element among hosts, or combinations of these processes. Rapid plasmid gains, losses, recombination events, and copy-number differences have been observed even among closely related isolates collected during a single hospital outbreak [63]. Consequently, resistance-gene abundance cannot be interpreted in the same manner as the concentration of a conventional chemical pollutant: biological context, host association, mobility, and gene expression all influence its relevance [64]. Our multimodal design represents one approach to separating host association from PDS abundance in an antimicrobial-exposed microbiome.

The multimodal design was useful because each sequencing layer supplied a different form of evidence. Bulk short reads provided longitudinal quantitative depth, long reads improved recovery of longer PDS contigs, and single-cell reads provided cellular host context. PDS contigs preserved sequence provenance and ARG localization, whereas PDS bins reduced fragmentation by collapsing related sequences. The single-cell experiment was designed to maximize the number of cells surveyed rather than to maximize genome completeness per cell. Many semi-permeable capsules (SPCs) were therefore sequenced at comparatively low depth, which limited complete taxonomic characterization and reconstruction of plasmid-derived sequences. This reflects a study-specific balance between cell throughput and per-cell depth rather than an intrinsic limitation of the SPC platform, which can instead be configured with fewer cells and greater sequencing depth [14]. Nevertheless, the single-cell layer supported host attribution for a subset of ARG-bearing PDS contigs and bins. This extends recent work on single-cell and probe-based plasmid-host linkage by connecting host-resolved PDS evidence to longitudinal bulk metagenomic quantification [14], [15], [61].

Several PDS bins carried ribosomal-target resistance markers, but *cfrE* had the clearest direct relationship to pleuromutilin resistance [54], [55], [56]. *ermB* was interpreted more broadly as a 23S-rRNA-target resistance determinant rather than a tiamulin-specific marker [57]. Reduced tiamulin susceptibility can also arise through mutations in 23S rRNA and the ribosomal proteins L3 and L4, encoded by *rplC* and *rplD*, respectively [65], [66]. These chromosomal markers are distinct from PDS-associated ARG carriage, and the PDS catalogue therefore represents only one component of the possible tiamulin-resistance landscape. Although all three *cfrE* -bearing PDS bins were predicted as mobilizable, their limited representation and the absence of a host-resolved abundance response do not support identifying *cfrE* as a treatment-response driver in this cohort.

The exploratory response models support this cautious interpretation. Two PDS bins showed prevalence associations with the post-event period and/or the post-event-by-pen-side interaction, but neither association was host-resolved. No significant response was detected for PDS-bin abundance, PDS-associated ARG-family abundance, or host-attributed PDS-contig abundance. These findings do not reduce the value of the host-linked catalogue; rather, they show that the unplanned and confounded exposure event was not suitable for establishing a host-specific plasmid-mediated treatment response. The response analysis should therefore be interpreted as a stress test of the workflow and a means of prioritizing candidate PDS features for future investigation.

The principal limitations arose from attrition and taxonomic skew in the single-cell workflow, incomplete PDS reconstruction, and the unplanned treatment structure. The laboratory workflow targeted approximately 10,000 cells per sample at *λ* = 0.1 cells per SPC, meaning that most SPCs were expected to be empty. Additional losses occurred during recovery of the four-component barcode, low-read filtering, assembly, and SAG quality filtering. Of 16,512 evaluated computationally, 6,533 passed the catalogue-level SAG criteria. Some discarded SPCs may nevertheless have contained biologically valid cells that failed one or more conservative barcode-quality criteria. Conversely, some low-read products may have represented empty capsules or extracellular DNA that underwent amplification.

The high number of samples pooled into each barcoding run may also have reduced recovery relative to the earlier validation design, in which fewer samples were processed together [14]. The demultiplexing workflow had not been optimized to maximize recovery under this degree of multiplexing, leaving room for improvement in future studies. At the same time, the experiment deliberately favoured surveying more cells at lower per-cell sequencing depth rather than recovering nearly complete genomes from fewer cells. These design choices limited SAG completeness, taxonomic assignment, and PDS reconstruction.

Taxonomic recovery was dominated by a restricted set of predominantly Gram-positive taxa, potentially because the sonication-based detachment step damaged more fragile cells before encapsulation. Many bulk-derived taxa consequently lacked matching SAG evidence, and many PDS features lacked sufficient unique sequence or single-cell coverage for confident host linkage. No PDS feature predicted as conjugative was recovered in the retained single-cell data. Finally, the six pigs came from one farm, and tiamulin exposure was completely confounded with pen side and disease signs. These limitations restrict the biological interpretation to candidate host associations and workflow feasibility rather than causal treatment effects.

In conclusion, integrating longitudinal bulk metagenomics with long-read and single-cell evidence produced a host-resolved view of plasmid-derived resistance reservoirs in the pig gut microbiome. The workflow linked a conservative subset of ARG-bearing PDS bins and contigs to bacterial host taxa, while the unplanned tiamulin event produced only broad prevalence associations and no host-resolved abundance response. Future applications should optimize barcode recovery in highly multiplexed SPC runs, increase per-cell sequencing depth where complete genome or plasmid reconstruction is required, and evaluate planned antimicrobial perturbations with balanced replication. These improvements would strengthen the use of single-cell-informed PDS linkage for determining how mobile resistance features are distributed and change across bacterial host backgrounds.

## Acknowledgements

We thank the participating farmer and farm staff for enabling longitudinal sample collection. We also thank colleagues involved in sample handling, sequencing, and technical discussions.

Large language model tools were used for language editing, consistency checks, and software troubleshooting suggestions. The authors designed the study, performed the analyses, interpreted the results, wrote the manuscript, and verified all AI-assisted outputs.

## Data Availability

The sequence reads generated in this study have been deposited in the European Nucleotide Archive under accession number PRJEB120797. Analysis scripts, workflow documentation, and table-generation notebooks are available at https://github.com/zuvale/pigpen.git.

