## Supplementary Information for "Single-Cell Metagenomics Links Plasmid-Derived Antimicrobial Resistance to Hosts in the Pig Gut Microbiome"

#### Methods

##### Tool Versions

**Supplementary Table S2. Software versions and database releases used in this study.** Software versions were obtained from the analysis executables and recorded computational environments.

| Software | Version |
|---|---|
| Python | 3.11.14 |
| POD5 tools | 0.3.15 |
| Trim Galore! | 0.6.10 |
| MEGAHIT | 1.2.9 |
| Dorado | 0.9.6 |
| metaFlye (Flye) | 2.9.5 |
| Medaka | 2.0.1 |
| SPAdes | 4.1.0 |
| MetaBAT2 | 2.17 |
| MaxBin2 | 2.2.7 |
| CONCOCT | 1.1.0 |
| VAMB | 5.0.4 |
| LorBin | 0.1.0 |
| DAS Tool | 1.1.7 |
| CheckM2 | 1.1.0 |
| GUNC | 1.0.6 |
| dRep | 3.6.2 |
| GTDB-Tk | 2.4.1 |
| Prokka | 1.13 |
| MMseqs2 | 18.8cc5c |
| eggNOG-mapper | 2.1.12 |
| RGI | 6.0.5 |

| | |
|---|---|
| CoverM | 0.7.0 |
| featureCounts | 2.1.1 |
| SnapATAC2 | 2.4.0 |
| geNomad | 1.11.1 |
| SCAPP | 0.1.4 |
| MOB-Typer | 3.1.9 |
| Pling | 3.1.9 |
| CD-HIT | 4.8.1 |
| BWA-MEM2 | 2.2.1 |
| Mosdepth | 0.3.10 |
| MUMmer4/NUCmer | 4.0.0-beta2 |
| SciPy | 1.17.1 |

### MAG and SAG Catalogue Construction

Assembly and binning were performed at three sample granularities. In the single-sample mode, one sample was used to generate one assembly and genome collection. In the within-pig co-assembly mode, all samples from one pig were combined to generate one assembly and genome collection. In the whole-study co-assembly mode, all samples across the study were combined to generate one assembly and genome collection.

For MetaBAT2, MaxBin2, and CONCOCT, reads were mapped to assemblies according to the assembly granularity. In the single-sample mode, one sequence run was aligned to the corresponding assembly. In the within-pig co-assembly mode, all read pairs from the corresponding pig were aligned. In the whole-study co-assembly mode, all available read pairs were aligned. VAMB and LorBin were handled separately because they perform co-binning internally. For VAMB and LorBin, single-sample assemblies within a pig were concatenated for the single-sample strategy, and all within-pig co-assemblies were concatenated for the within-pig strategy.

Genome candidates from the different binners were refined with DAS Tool within each sample-granularity layer. Refined genomes were assessed with CheckM2 and GUNC. MAGs were retained if they had at least 75% completeness, at least 80% coding density, and no more than 10% contamination. SAGs were retained if they had at least 20% completeness, no more than 30% contamination, and a clade separation score below 0.5. MAGs were dereplicated with dRep using 95% primary and 99% secondary ANI thresholds. SAGs were not dereplicated because they represented single-cell-derived assemblies.

Taxonomic annotation of MAGs and SAGs was performed with GTDB-Tk. Coding sequences were predicted with Prokka. Protein sequences from MAGs and SAGs were clustered with MMseqs2 at 90% amino-acid identity to generate a non-redundant gene catalogue. Gene clusters were annotated with eggNOG-mapper and RGI. Bulk metagenomes and gene features were quantified with CoverM and featureCounts.

### Single-Cell Barcode Demultiplexing and SPC Recovery

Raw reads were assigned to SPCs using the four-component combinatorial barcode scheme provided by Atrandi Biosciences. Read sets with too few read pairs were excluded due to low-read products being considered more likely to represent empty or weakly amplified SPCs. Following demultiplexing and read filtering, 16,512 read sets were assembled individually and evaluated as candidate SAGs. The number of retained SPCs was substantially lower than the nominal number loaded into the workflow. Most loaded SPCs were expected to be empty because the target occupancy was lambda = 0.1 cells per SPC. Further loss occurred when barcode components failed quality control, when demultiplexed read sets did not meet the minimum read threshold, or when assemblies did not meet the SAG quality criteria. Consequently, the final SAG catalogue represents a conservative subset of the cell-containing SPCs that entered sequencing.

### Single-cell Additional Filtering for Functional Embedding

Single-cell barcodes were retained as cells if they passed genome-quality and feature-content filters. Barcodes were retained if they had at least 45% completeness, at least 800 annotated genes, and genes assigned to at least 0.4% of the final non-redundant gene catalogue.

Gene-feature filtering followed single-cell count-preprocessing logic by retaining informative features while trimming both tails of the summed-count distribution. A gene feature was retained if it was detected in at least seven cells, or if its log1p-transformed summed featureCounts read count was greater than 3 and less than 7. The top 200,000 retained features were used for dimensionality reduction. Functional count matrices were embedded using Laplacian eigenmaps, a k-nearest-neighbour graph was constructed, Leiden clustering was performed at resolution 1, and UMAP was used for two-dimensional visualization.

### PDS Recovery, Annotation, Quantification, and Host Linkage

#### Candidate PDS Discovery

Candidate plasmid-derived sequences were recovered from four discovery routes: geNomad predictions from Illumina bulk assemblies, geNomad predictions from ONT bulk assemblies, circular metaFlye ONT contigs using Medaka-polished consensus sequences downstream, and SCAPP predictions from single-cell assemblies. Candidate identifiers were standardized to encode sequencing technology, prediction tool, sample, length, and sequence hash before downstream catalogue construction.

#### Candidate Dereplication And Confidence Filtering

Candidate sequences were pooled and dereplicated in several stages. Exact duplicate sequences were removed first. Final candidate clustering used MMseqs2 with minimum sequence identity 0.95 and coverage 0.90. MOB-Typer was then applied to annotate plasmid marker genes and nearest-neighbour information.

Candidate confidence classes were defined from similarity to known plasmids and plasmid marker-gene evidence. Very high confidence required MASH nearest-neighbour distance <= 0.06. High confidence required MASH nearest-neighbour distance > 0.06 and either an annotated replicon type or at least two of relaxase, mating-pair formation, and oriT annotations. Middle confidence required MASH nearest-neighbour distance > 0.06 and one of replicon, relaxase, mating-pair formation, or oriT annotation. Low confidence was assigned otherwise. Low-confidence sequences and sequences shorter than 1,000 bp were discarded. The retained middle-, high-, and very high-confidence sequences constituted the PDS-contig catalogue.

#### PDS Binning With Pling

Retained PDS contigs were clustered into PDS bins with Pling. Pling was run with containment distance 0.3 and DCJ thresholds of 4, 3, and 2. The final bin catalogue used the dcj_thresh_2_graph output, and bins were defined using the full Pling type/subcommunity grouping. Sequence-to-bin mappings were retained so that PLS contigs could be summarized at the PLB-bin level while preserving sequence-level provenance.

Within-bin containment filtering was used to identify shared and informative PDS regions. Filtering used 99% ANI, minimum alignment length 500 bp, drop-covered fraction 0.95, minimum unique length 500 bp, and minimum unique fraction 0.05. PDS bins are therefore collapsed groups of related retained PDS contigs, not necessarily complete biological plasmids.

#### PDS Annotation

PDS contigs and PDS bins were annotated separately. PDS-contig annotation used Prokka-predicted coding sequences from retained PDS contigs and ran eggNOG-mapper and RGI on the contig-level annotation layer. PDS-bin annotation used Prokka-predicted coding sequences from bin-level sequence outputs. For PDS bins, translated coding sequences were redundancy-reduced with CD-HIT before eggNOG-mapper and RGI annotation. RGI-derived ARG annotations were categorized into resistance-mechanism groups and written separately for PDS-bin and PDS-contig analysis levels.

The bin-level ARG table contains PDS-bin identifiers, ARG labels, ARG categories, and predicted mobility classes. It was used for collapsed catalogue summaries and ARG co-occurrence analyses. The contig-level ARG table contains PDS-bin identifiers, PDS-contig identifiers, ARG labels, ARG categories, and predicted mobility classes. It was used for ARG localization and host-resolved inspection. Keeping these two tables separate prevents collapsed bin-level annotations from being mixed with contig-level sequence evidence.

#### Informative PDS Regions and Bulk Quantification

For PDS quantification, retained PDS sequences were aligned against themselves with MUMmer/NUCmer to identify shared sequence. Shared regions required at least 95% identity and at least 100 bp aligned. For PDS-contig-level analyses, sequence shared between retained PDS contigs was masked. For PDS-bin-level analyses, alignments between PDS contigs belonging to the same PDS bin were ignored, and only sequence shared between different PDS bins was masked. Regions shared with MAG sequences were also masked for bulk quantification. Informative regions shorter than 500 bp were excluded.

Bulk short reads were aligned to a combined PDS/MAG reference with BWA-MEM2. PDS depth was summarized over informative regions using Mosdepth-derived regional coverage. PDS-contig and PDS-bin depth summaries were generated separately. Bin-level summaries aggregated informative regions according to the sequence-to-bin mapping. Depth summaries included median depth, mean depth, trimmed mean depth, maximum depth, informative-region length, informative fraction, and depth-profile quality flags.

Genome-normalized abundance was calculated as PDS depth divided by host species marker depth. Host marker depth was summarized from single-copy marker depths. Host marker depth filters required at least 10 markers, at least 50% nonzero markers, and median marker depth of at least 1. Host-depth cutoffs were host depth >= 2.0 for the permissive linkage layer and host depth >= 5.0 for the stringent linkage layer. Because PDS recovery is often incomplete and quantification uses informative recovered regions, genome-normalized abundance was interpreted as a recovered-region abundance estimate rather than literal complete-plasmid copy number.

#### Single-cell PDS Evidence

Single-cell reads were aligned to retained PDS sequences with BWA-MEM2, and barcode-level coverage over informative PDS regions was calculated with Mosdepth. For single-cell quantification, regions shared between PDS features and MAG/SAG assemblies were identified by PDS-to-genome alignment. In addition, for each barcode, regions shared between a PDS feature and that barcode's own SAG assembly were subtracted before quantification. This made barcode-level PDS evidence depend on sequence not already represented in the assembled SAG. Quantification was performed separately for PDS-bin and PDS-contig evidence layers.

Evidence for a PDS feature in a barcode was classified as strong if breadth of coverage at 1x was >= 0.30 or breadth of coverage at 3x was >= 0.20. Weak evidence was defined as breadth of coverage at 1x >= 0.20 if the observation was not already strong. Features with no retained informative region were excluded from the corresponding evidence layer.

#### Host-Linkage Layers

Barcode-level PDS evidence was joined to taxonomically assigned cells to create host-support tables. Host support was computed separately for PDS bins and PDS contigs. For each PDS-feature/species pair, we recorded the number of supporting cells and compared observed support to the background distribution of that PDS feature across taxonomically assigned cells using a one-sided Fisher exact test. P-values were corrected with the Benjamini-Hochberg method.

Stringent host assignments required at least two supporting cells and q < 0.05. The permissive layer retained all observed host-feature pairs from the support table, including one-cell links and links that did not pass correction. Stringent linkage was used for conservative host-linkage claims and visual summaries. Permissive linkage was retained for sensitivity analyses and exploratory abundance modelling. PDS-bin linkage was used as the primary collapsed host-linkage level; PDS-contig linkage was used for ARG localization, host-resolved inspection, and sensitivity analyses.

#### Mobilome Annotation

MAGs, SAGs, and PDS sequences were further annotated with general mobile genetic elements using the EBI Mobilome Annotation Pipeline [1]. For PDS sequences, sequences below 1,000 bp were excluded because of a pipeline-internal length check. After MGE prediction, coding sequences from the MAG, SAG, and PDS catalogues were annotated according to whether they partially or fully overlapped with a predicted non-plasmidic mobile genetic element. This analysis was treated as exploratory and was not used for primary host-PDS claims.

#### Differential Response Testing

We used MaAsLin3 [s2] to screen for associations between abundance or prevalence features and the tiamulin exposure event. The tested feature groups were metagenome-assembled genome relative abundance, order-attributed antimicrobial resistance gene abundance, microbiome-wide PDS-bin abundance, PDS-associated ARG-family abundance, and host-attributed PDS-contig genome-normalized abundance. PDS-bin models were treated as the primary collapsed PDS response screen. PDS-contig models were treated as exploratory sensitivity analyses because contigs preserve ARG localization but are more fragmented and more numerous than collapsed PDS bins.

For the event-response screen, the model formula in R mixed-effect syntax was:

~ ordered(week) + post_event * pen_side + (1|pig)

Here, week represents ordered sampling time, post_event distinguishes samples collected before the end of antimicrobial treatment from those collected after treatment, and pen_side distinguishes the left-side reference pigs B, D, and F from the right-side exposed pigs A, C, and E. Ordered week was included to account for gradual longitudinal change, whereas post_event tested for an additional step-like shift after the treatment boundary.

The interaction term was interpreted as a difference-in-differences-like contrast, asking whether the post-event change in the exposed right-side pigs exceeded the corresponding temporal change in the left-side reference pigs. Because treatment was unplanned and confounded with pen side and the observed diarrhoea event, all response models were interpreted as exploratory screening analyses rather than fully powered causal treatment-response models.

### Results

#### PDS Catalogue Composition

The supplementary catalogue tables provide the source counts underlying the main PDS recovery result. They summarize the number of input sequencing files and assemblies, the retained MAG and SAG catalogues, the retained PDS sequence catalogue, PDS-bin composition, technology support, confidence classes, and mobility classes. These tables support the main-text catalogue summary without repeating all values in the Results.

**Supplementary Table S3. Data inputs and PDS catalogue composition.**

Summary of sequencing inputs, genome catalogues, retained PDS sequences, PDS bins, host-attributed PDS bins, bin composition, technology support, confidence classes, and predicted mobility classes. Values combine the data-input summary and PDS-binning catalogue summary.

**S3A. Data and catalogue output summary**

| Result | Unit | Value |
|---|---|---|
| Bulk metagenomes in longitudinal metadata | samples | 64 |
| Raw Illumina read files | files | 128 |
| Raw Illumina read pairs | read pairs | 1,767,648,350 |
| Nanopore pig-level POD5 runs | runs | 6 |
| Nanopore classified barcode FASTQs | files | 18 |
| SAG assemblies assessed before QC | assemblies | 16,512 |
| SAG assemblies retained after filtering | cells | 6,533 |
| Taxonomically labelled SAGs | cells | 6,532 |
| MAGs in dereplicated catalogue | genomes | 1,181 |
| Retained PDS sequences after filtering | sequences | 885 |
| PDS bins after collapsing related sequences | bins | 195 |
| ARG-bearing PDS bins | bins | 28 |
| Stringent host-attributed PDS bins | bins | 11 |
| Stringent host-PDS bin pairs | pairs | 17 |

**S3B. PDS binning and catalogue composition**

| Summary | Value |
|---|---|
| Retained PDS sequences | 885 |
| PDS bins | 195 |
| Singleton PDS bins | 128 |
| Multi-sequence PDS bins | 67 |
| Largest PDS bin | PLB_000026, 152 retained sequences |

| | |
|---|---|
| Bins supported by >1 sequencing technology | 33 |
| Bins supported by all three sequencing technologies | 4 |
| Bin confidence distribution | Middle: 71; High: 53; Very High: 71 |
| Bin mobility distribution | Non-mobilizable: 116; mobilizable: 77; conjugative: 2 |

#### ARG Annotation

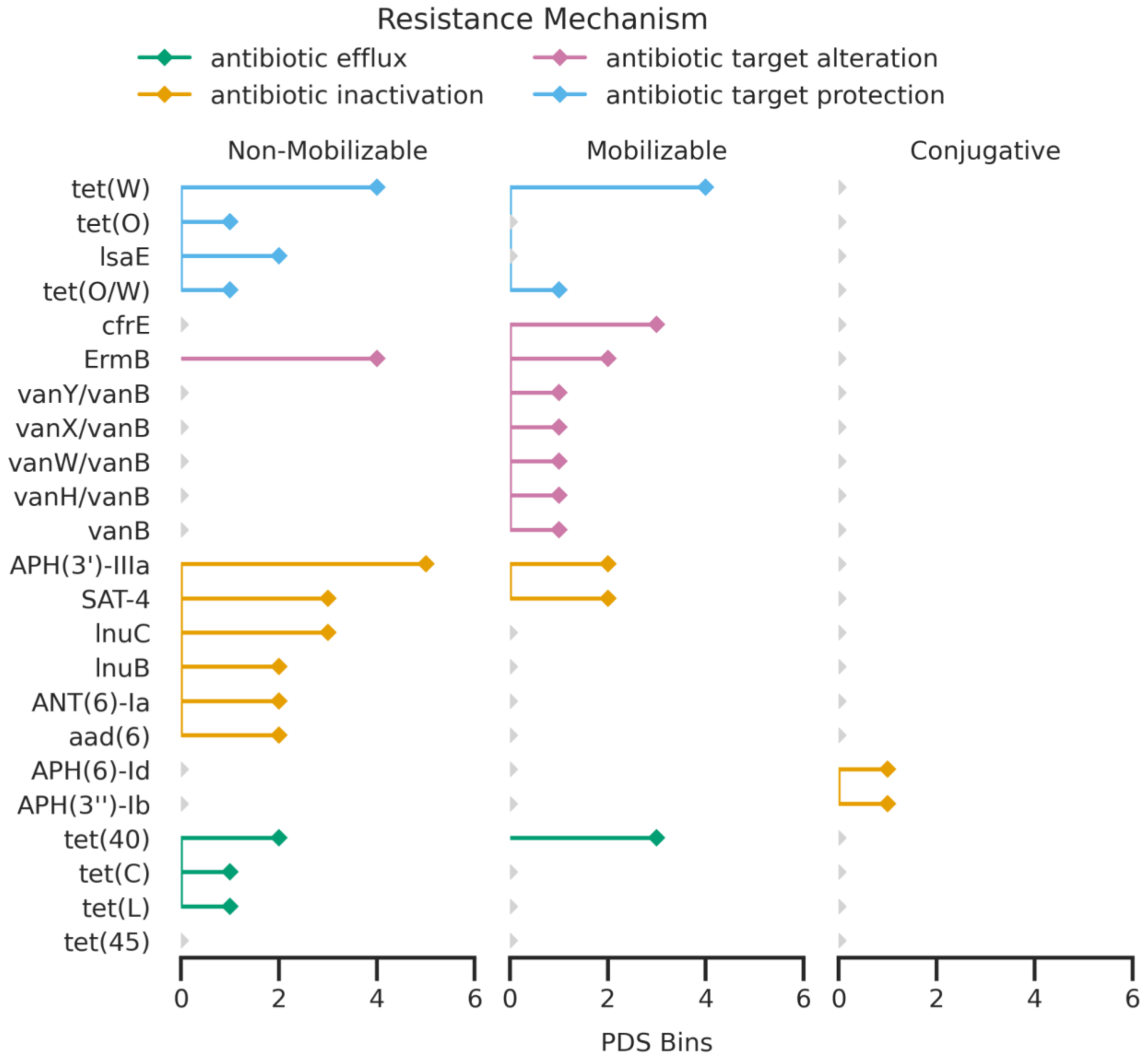


**Figure S1. Bin-level ARG content of the PDS catalogue.**
Number of ARG-bearing PDS bins carrying each ARG annotation, stratified by predicted PDS-bin mobility class and coloured by resistance mechanism. This figure provides the bin-level companion to the main contig-level ARG summary in Figure 3A. Grey markers indicate ARG labels not observed in the corresponding mobility class.

--------------------------------------------------------------------------------------------------------------------

ARG annotation was summarized separately at PDS-contig and PDS-bin levels. The contig-level summary preserves sequence-level localization and supports host-resolved inspection in the main text. The bin-level summary collapses related PDS contigs and was used for catalogue-level ARG counts and co-occurrence analyses.

**Supplementary Table S4. ARG-bearing PDS bins by resistance label and predicted mobility class.**

Number of PDS bins carrying each of the 22 RGI-derived ARG labels, stratified by predicted PDS-bin mobility class and resistance mechanism.

| Best hit ARO | Resistance mechanism | PDS bins | Non-mobilizable | Mobilizable | Conjugative |
|---|---|---|---|---|---|
| tet(W) | antibiotic target protection | 8 | 4 | 4 | 0 |
| APH(3')-IIIa | antibiotic inactivation | 7 | 5 | 2 | 0 |
| ErmB | antibiotic target alteration | 6 | 4 | 2 | 0 |
| SAT-4 | antibiotic inactivation | 5 | 3 | 2 | 0 |
| tet(40) | antibiotic efflux | 5 | 2 | 3 | 0 |
| cfrE | antibiotic target alteration | 3 | 0 | 3 | 0 |
| lnuC | antibiotic inactivation | 3 | 3 | 0 | 0 |
| ANT(6)-Ia | antibiotic inactivation | 2 | 2 | 0 | 0 |
| aad(6) | antibiotic inactivation | 2 | 2 | 0 | 0 |
| lnuB | antibiotic inactivation | 2 | 2 | 0 | 0 |
| lsaE | antibiotic target protection | 2 | 2 | 0 | 0 |
| tet(O/W) | antibiotic target protection | 2 | 1 | 1 | 0 |
| APH(3'')-Ib | antibiotic inactivation | 1 | 0 | 0 | 1 |
| APH(6)-Id | antibiotic inactivation | 1 | 0 | 0 | 1 |
| tet(C) | antibiotic efflux | 1 | 1 | 0 | 0 |

| | | | | | |
|---|---|---|---|---|---|
| tet(L) | antibiotic efflux | 1 | 1 | 0 | 0 |
| tet(O) | antibiotic target protection | 1 | 1 | 0 | 0 |
| vanB | antibiotic target alteration | 1 | 0 | 1 | 0 |
| vanH/vanB | antibiotic target alteration | 1 | 0 | 1 | 0 |
| vanW/vanB | antibiotic target alteration | 1 | 0 | 1 | 0 |
| vanX/vanB | antibiotic target alteration | 1 | 0 | 1 | 0 |
| vanY/vanB | antibiotic target alteration | 1 | 0 | 1 | 0 |

### Single-Cell PDS-Bin Evidence

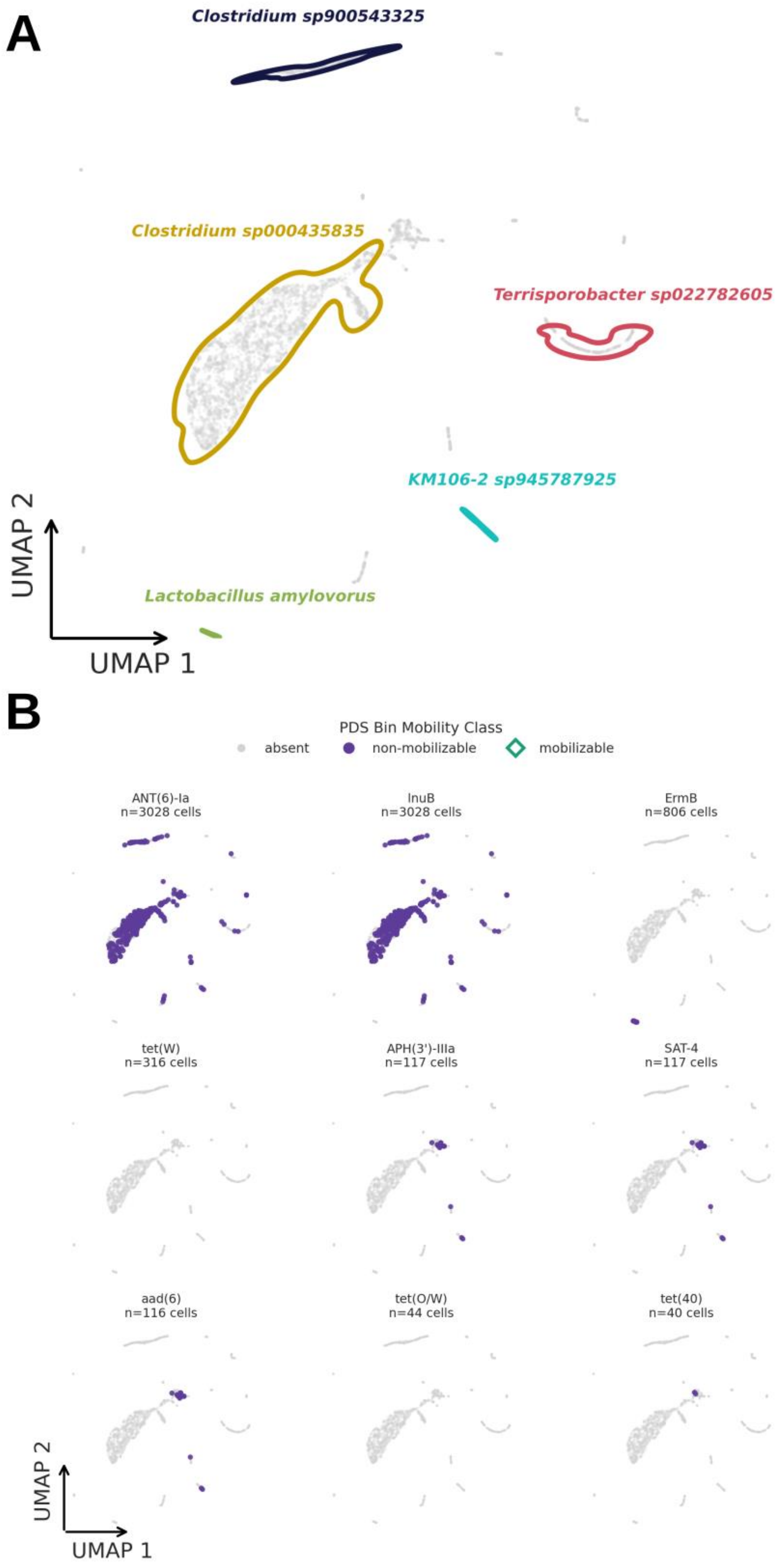
A
Clostridium sp900543325
Clostridium sp000435835
Terrisporobacter sp022782605
KM106-2 sp945787925
Lactobacillus amylovorus
UMAP 2
UMAP 1
B
PDS Bin Mobility Class
absent
non-mobilizable
mobilizable
ANT(6)-Ia
n=3028 cells
lnuB
n=3028 cells
ErmB
n=806 cells
tet(W)
n=316 cells
APH(3')-IIIa
n=117 cells
SAT-4
n=117 cells
aad(6)
n=116 cells
tet(O/W)
n=44 cells
tet(40)
n=40 cells
UMAP 2
UMAP 1

**Figure S2. Single-cell evidence for host-associated ARG-bearing PDS bins.**
(A) UMAP of functional embeddings derived from the filtered SAG catalogue. The embedding is based on functional annotations after filtering cells and functional features, and the five species with the highest cell counts are highlighted. (B) Per-cell evidence for ARG-bearing PDS bins, separated by ARG annotation and predicted PDS-bin mobility class. Grey points indicate cells without evidence for the displayed feature, filled points indicate evidence for non-mobilizable PDS bins, and open diamonds indicate evidence for mobilizable PDS bins.

----------------------------------------------------------------------------------------------------------------------

PDS-bin evidence provides a collapsed view of single-cell PDS detection. This layer is less resolved than the contig-level view used in the main text, but it is useful for assessing whether related PDS sequences show consistent barcode-level evidence across cells and taxa.

**Supplementary Table S5. Single-cell PDS-bin detection summary.**

Number of barcode-level PDS-bin observations, cells, PDS bins, and ARG-bearing PDS bins detected using strong and weak single-cell evidence thresholds. Breadth values summarize coverage over retained informative PDS-bin regions.

| **Evidence level** | **Observations** | **Cells** | **PDS bins** | **ARG-bearing PDS bins** | **Median breadth ≥1x** | **Median breadth ≥3x** |
|---|---|---|---|---|---|---|
| Strong | 9,584 | 4,216 | 92 | 10 | 0.455 | 0.386 |
| Weak | 2,151 | 1,619 | 64 | 9 | 0.231 | 0.164 |

### Host-Linkage and Bin-Level Networks

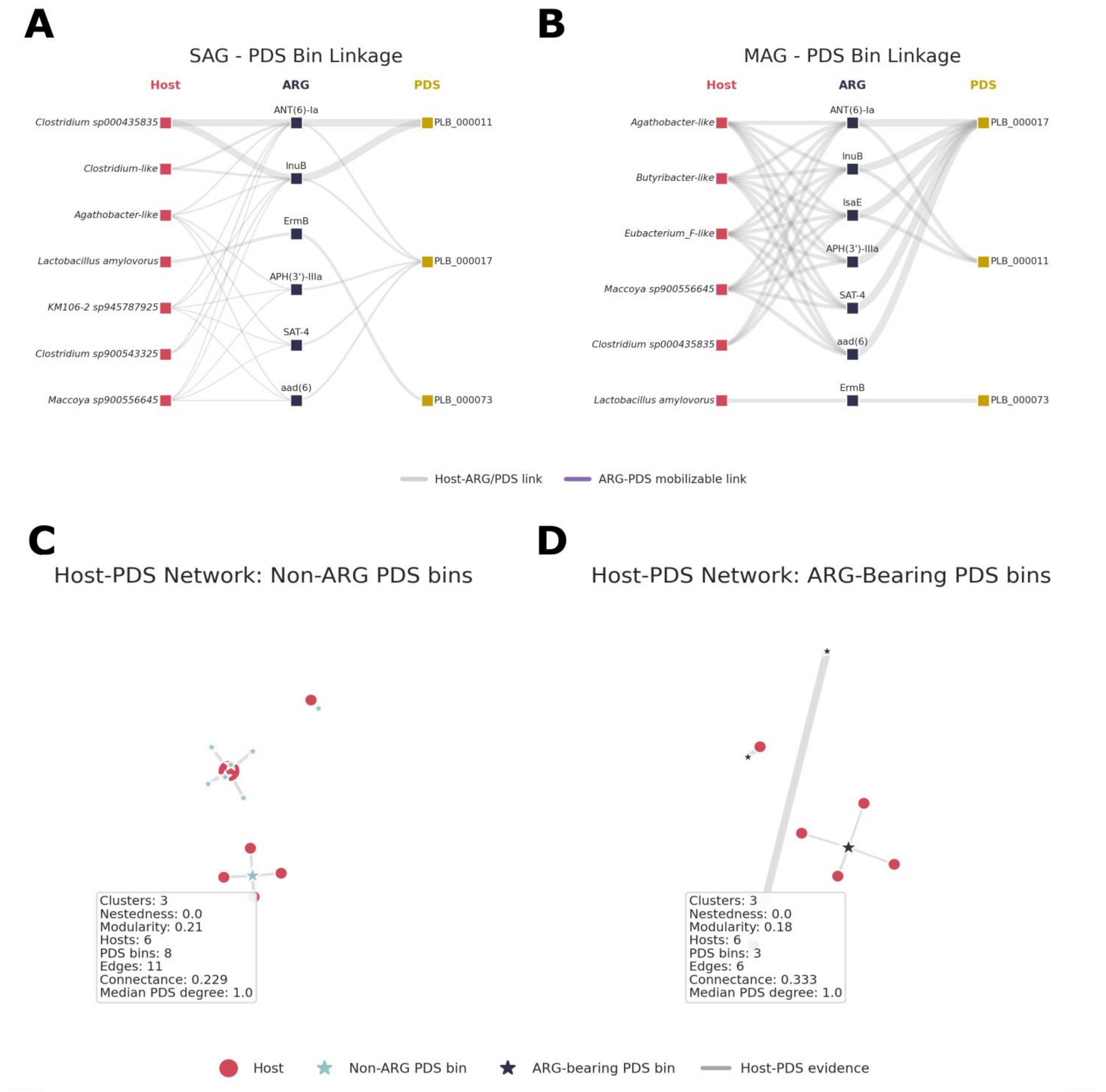


**Figure S3. Host attribution of ARG-bearing PDS bins.**
(A,B) Sankey-style diagrams showing associations between bacterial hosts, antimicrobial resistance genes, and PDS bins. Link thickness summarizes the number of unique host-ARG or ARG-PDS attributions. Purple ARG-PDS links indicate that the PDS bin is predicted as mobilizable. SAG-side plots use strong single-cell evidence for the displayed host-ARG-PDS triads, whereas MAG-side plots use stringent host support. (C,D) Bipartite host-PDS networks comparing non-ARG-bearing and ARG-bearing PDS bins. Network edges were derived from host-linkage layers and are shown as exploratory descriptors of host-linkage graph structure.

---

The bin-level linkage figures complement the main contig-level host-attribution results. PDS-bin displays reduce fragmentation by collapsing related PDS contigs, whereas PDS-contig displays retain sequence-level ARG localization. Network descriptors are reported as exploratory summaries of graph structure and should not be interpreted as direct measurements of transfer or biological host range.

**Supplementary Table S6. Display scope for host-ARG-PDS Sankey diagrams.**

Input and displayed scope for the host-ARG-PDS Sankey diagrams at SAG and MAG levels, shown separately for PDS-bin and PDS-contig evidence layers.

| Scope | Input units | Input hosts | Input ARGs | Input PDS | Input triads | Displayed hosts | Displayed ARGs | Displayed PDS | Displayed links |
|---|---|---|---|---|---|---|---|---|---|
| SAG PDS bin | 197 | 26 | 7 | 5 | 85 | 7 | 6 | 3 | 30 |
| MAG PDS bin | 6 | 6 | 7 | 3 | 28 | 6 | 7 | 3 | 38 |
| SAG PDS contig | 459 | 92 | 11 | 26 | 280 | 7 | 10 | 11 | 55 |
| MAG PDS contig | 37 | 21 | 11 | 13 | 73 | 10 | 10 | 10 | 68 |

**Supplementary Table S7. Host-linkage support by evidence scope.**

Summary of stringent and permissive host-linkage layers for PDS bins and PDS contigs. Stringent linkage required at least two supporting cells and FDR-corrected enrichment, whereas permissive linkage retained all observed host-feature pairs.

| Evidence scope | Filter | Host-feature pairs | Unique PDS features | Host species | ARG-bearing PDS features | ARG-bearing host-feature pairs | Median supporting cells | Maximum supporting cells |
|---|---|---|---|---|---|---|---|---|

| | | | | | | | | |
|---|---|---|---|---|---|---|---|---|
| PDS bin | stringent | 17 | 11 | 11 | 3 | 6 | 3.0 | 359 |
| PDS bin | permissive | 97 | 34 | 60 | 6 | 37 | 1.0 | 616 |
| PDS contig | stringent | 67 | 36 | 25 | 13 | 37 | 2.0 | 359 |
| PDS contig | permissive | 517 | 135 | 165 | 30 | 244 | 1.0 | 616 |

### Genome-Normalized Abundance

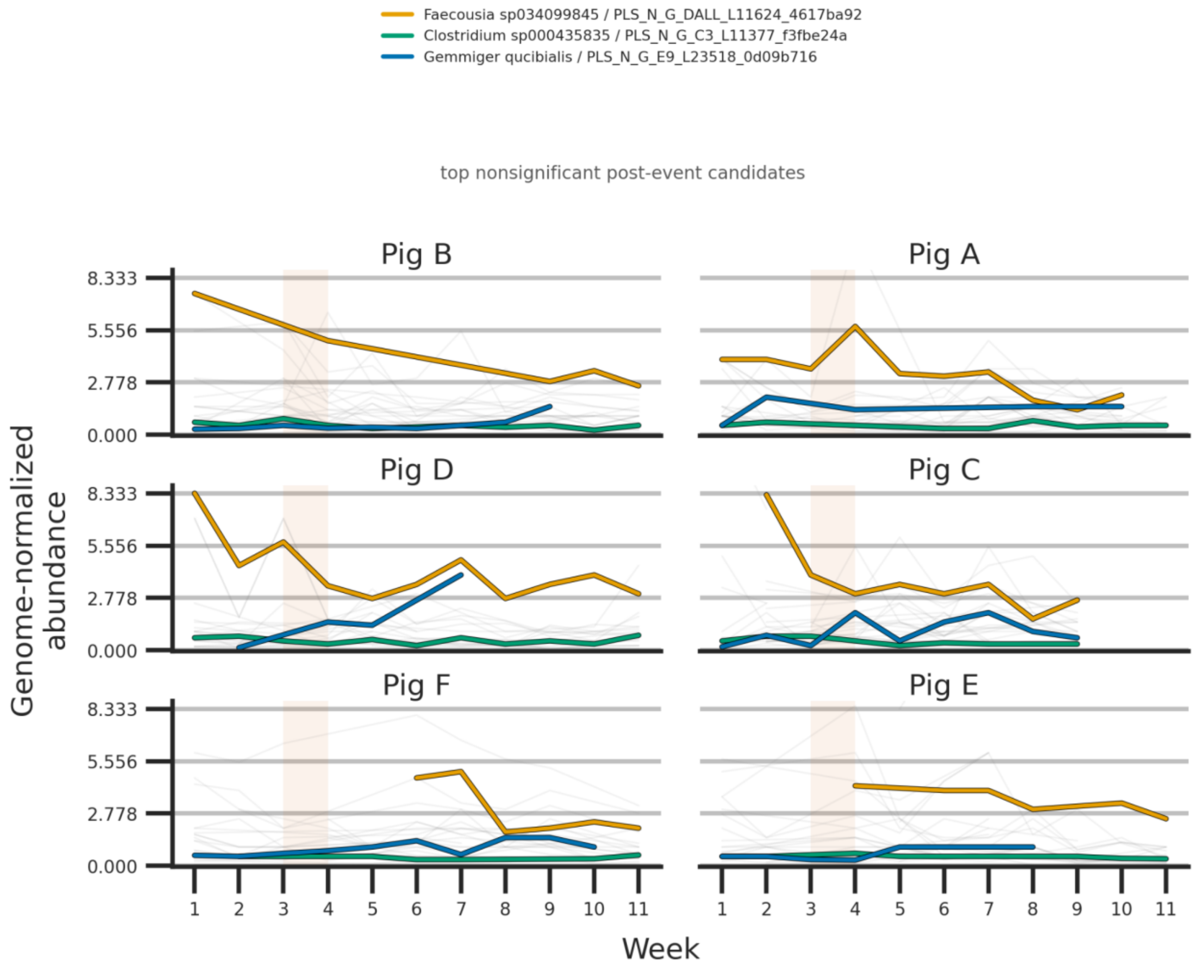


**Figure S4. Genome-normalized abundance trends for selected host-attributed PDS contigs.**
Longitudinal genome-normalized abundance estimates are shown for selected host-PDS-contig pairs across pigs and sampling weeks. Coloured lines highlight selected non-significant post-event candidate host-PDS-contig pairs, while pale background lines show additional retained host-PDS-contig observations. The shaded region marks the tiamulin exposure period. Values represent informative recovered PDS regions normalized by host marker depth and should not be interpreted as exact complete-plasmid copy number.

---

Genome-normalized abundance estimates were calculated over informative recovered PDS regions and normalized by host marker depth. These values are therefore used as abundance estimates for host-attributed PDS evidence, not as exact complete-plasmid copy numbers.

**Supplementary Table S8. Corrected genome-normalized PDS abundance summary.**

Summary of genome-normalized PDS abundance estimates after informative-region filtering and host-depth filtering. Values represent recovered informative PDS regions normalized by host marker depth and should not be interpreted as exact complete-plasmid copy number.

| Analysis unit | Observations | Samples | Pigs | PDS features | Host species | Median genome-normalized abundance | 95th percentile | Maximum |
|---|---|---|---|---|---|---|---|---|
| PDS bin, stringent host linkage | 14 | 14 | 6 | 1 | 1 | 0.400 | 0.820 | 0.857 |
| PDS bin, permissive host linkage | 421 | 63 | 6 | 7 | 12 | 0.774 | 3.500 | 10.500 |
| PDS contig, stringent host linkage | 126 | 35 | 6 | 6 | 3 | 0.789 | 2.554 | 4.000 |
| PDS contig, permissive host linkage | 1,161 | 64 | 6 | 21 | 21 | 1.000 | 4.500 | 27.000 |

### Broad Metagenome and PDS-ARG Response

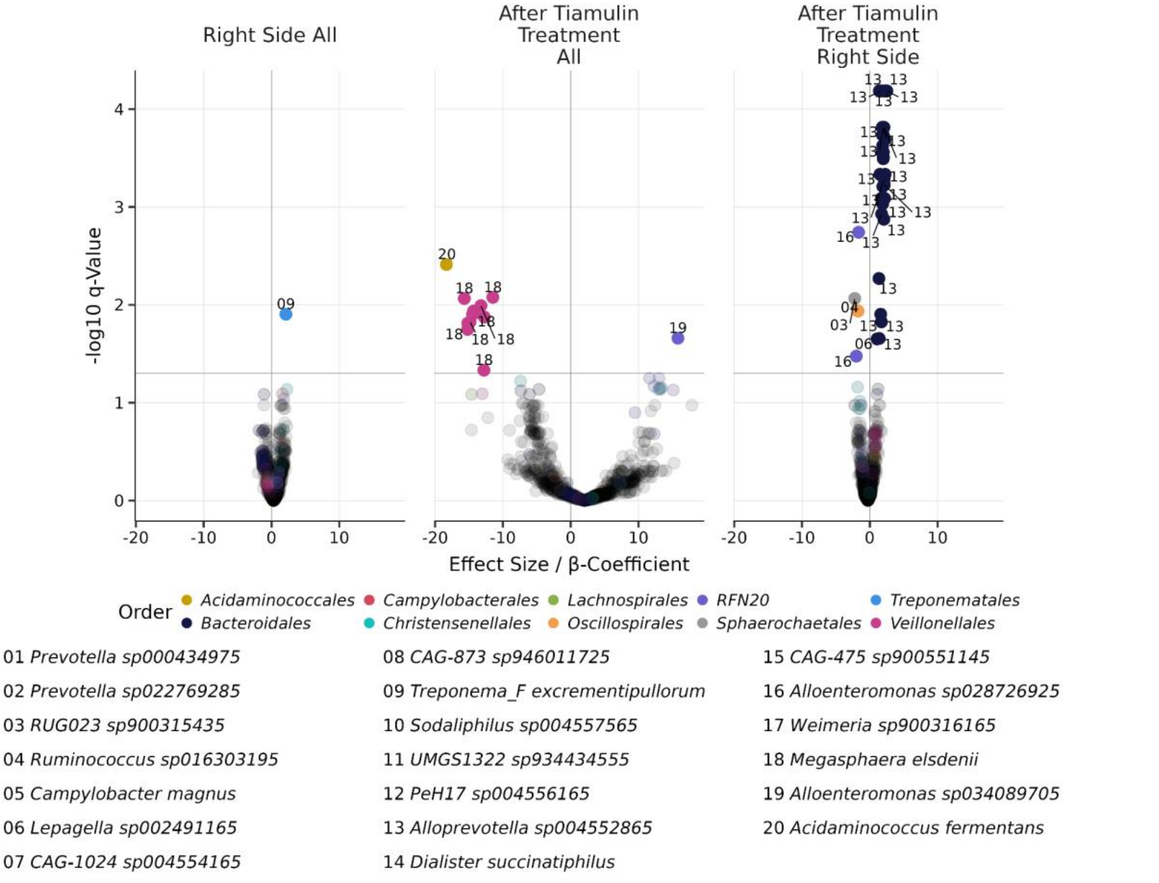


**Figure S5. Broad MAG-level abundance response screen in bulk metagenomes.** Volcano plots summarize MaAsLin3 abundance-model results for MAG-level relative abundance. Points are coloured by bacterial order where highlighted. Model terms represent right-side pen location, post-event period, and the post-event-by-right-side interaction. This broad MAG-level screen is shown as metagenomic background and is not a host-PDS analysis.

---

The response screens are included as exploratory background for the natural tiamulin exposure event. Broad MAG-level and order-attributed ARG models describe overall community and resistome context. PDS-bin, PDS-associated ARG-family, and host-attributed PDS-contig models address the plasmid-derived sequence catalogue more directly. Because treatment was unplanned and confounded with pen side, these models are interpreted as screening analyses rather than causal treatment-response tests.

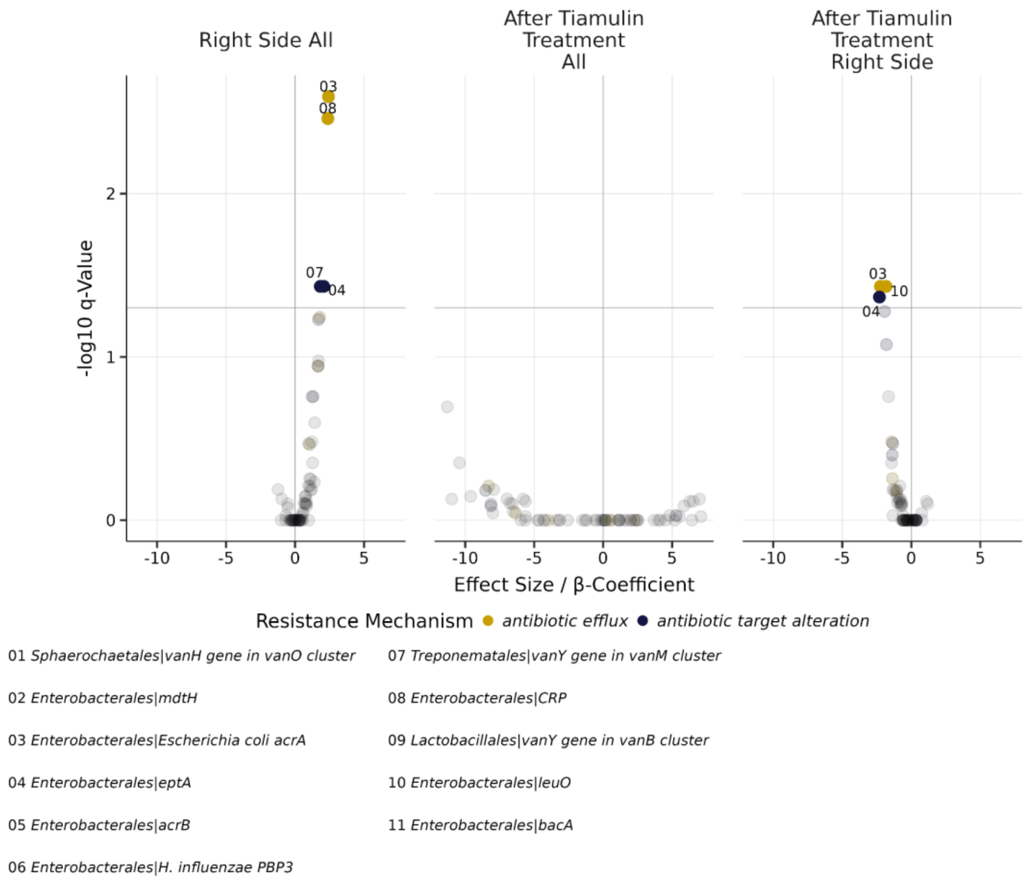


**Figure S6. Broad order-attributed ARG abundance response screen in bulk metagenomes.**
Volcano plots summarize MaAsLin3 abundance-model results for ARG features attributed to bacterial orders. Points are coloured by resistance mechanism where highlighted. Model terms represent right-side pen location, post-event period, and the post-event-by-right-side interaction. This screen provides broad resistome context and is not host-resolved at the PDS level.

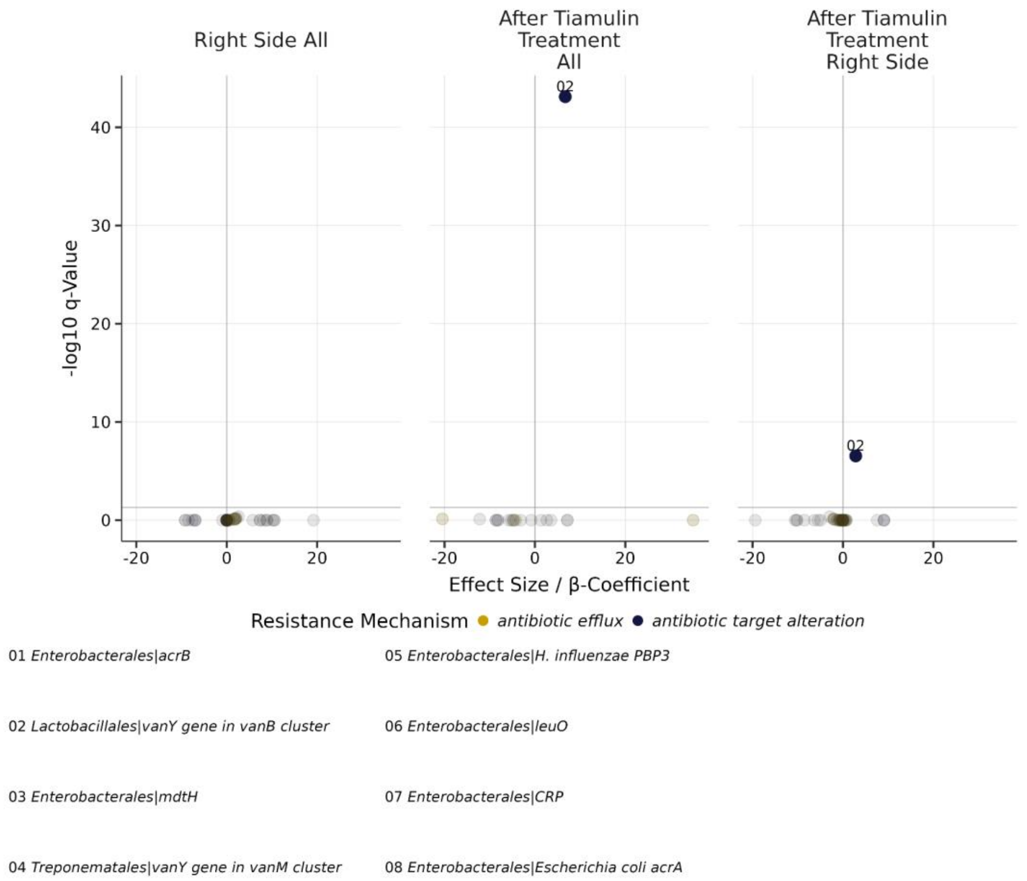


**Figure S7. Broad order-attributed ARG prevalence response screen in bulk metagenomes.**

Volcano plots summarize MaAsLin3 prevalence-model results for ARG features attributed to bacterial orders. Points are coloured by resistance mechanism where highlighted. Model terms represent right-side pen location, post-event period, and the post-event-by-right-side interaction. This screen provides broad resistome context and is not host-resolved at the PDS level.

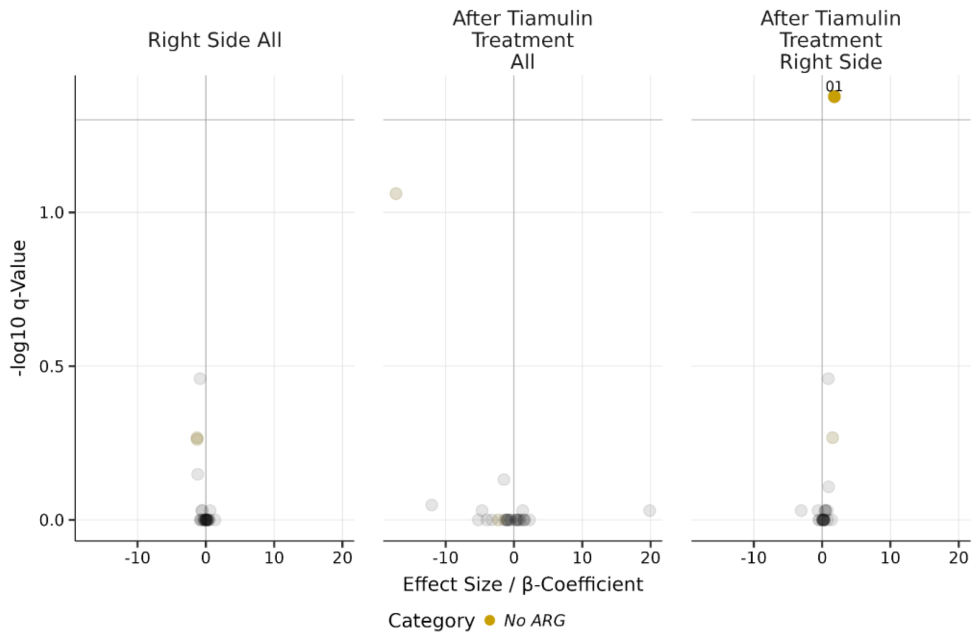


01 *PLB_000015* 02 *PLB_000190*

**Figure S8. Microbiome-wide PDS-bin abundance response screen.**
**V**olcano plots summarize component-specific, Benjamini–Hochberg-corrected q-values from MaAsLin3 abundance models of microbiome-wide PDS-bin abundance. Points represent PDS-bin features, and labelled points are selected top associations. One abundance interaction (PLB_000015; q = 0.042) met the component-specific FDR threshold; no abundance post-event main effect did. These exploratory associations are not causal treatment-response evidence because exposure was unplanned and confounded with pen side and disease signs.

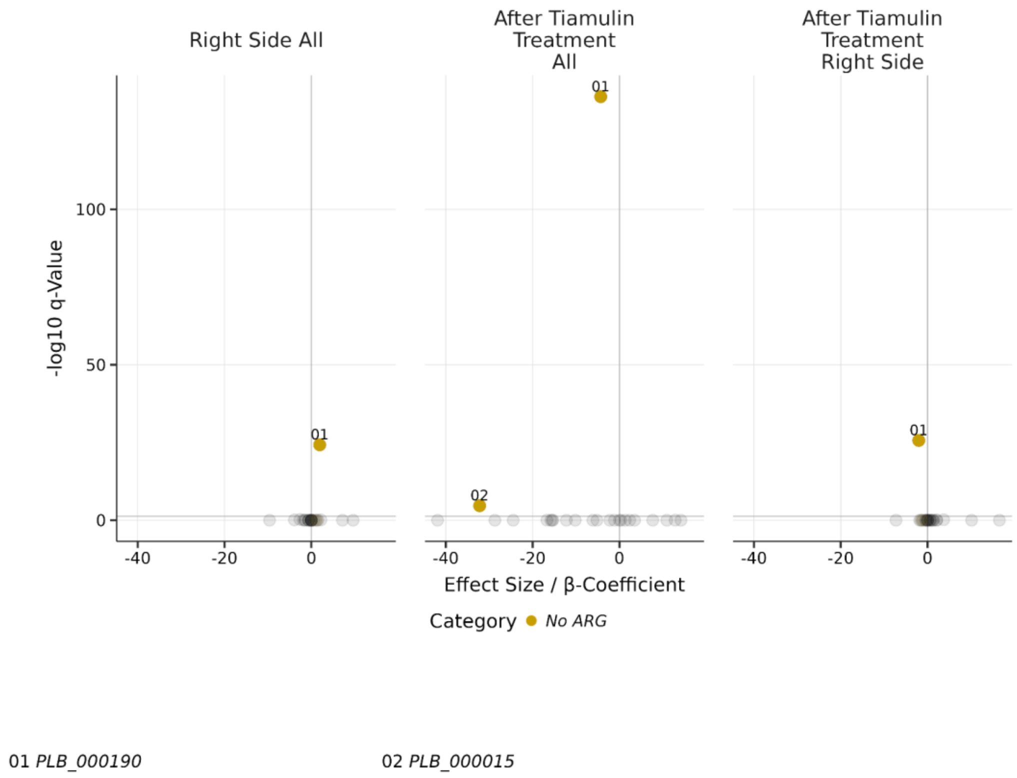


**Figure S9. Microbiome-wide PDS-bin prevalence response screen.**
Volcano plots summarize component-specific, Benjamini–Hochberg-corrected q-values from MaAsLin3 prevalence models of microbiome-wide PDS-bin prevalence. These prevalence components contribute to the two feature-level joint associations summarised in Supplementary Table S9. This screen complements the PDS-bin abundance model and is interpreted as exploratory rather than causal treatment-response evidence.

**Supplementary Table S9. Corrected MaAsLin3 model summary.**
Summary of exploratory MaAsLin3 model results. For the PDS-bin abundance/prevalence screen, reported q-values are feature-level joint q-values across abundance and prevalence components; "Top model component" identifies the component contributing the minimum q-value. A feature was called significant only when q < 0.05 and |β| > 0.5.

| **Model** | **Term** | **Tested features** | **Significant features** | **Minimum q value** | **Top feature** | **Top model component** |
|---|---|---|---|---|---|---|
| PDS-bin abundance/prevalence | post-event × pen-side interaction | 28 | 2 | 2.42e-26 | PLB_000190 | prevalence |

| | | | | | | |
|---|---|---|---|---|---|---|
| PDS-bin abundance/prevalence | post-event main effect | 29 | 2 | 6.39e-137 | PLB_000190 | prevalence |
| ARG-family PDS abundance | post-event × pen-side interaction | 11 | 0 | 0.403 | APH(3')-IIIa | abundance |
| ARG-family PDS abundance | post-event main effect | 11 | 0 | 0.935 | tet(40) | abundance |
| Host-PDS-contig abundance | post-event × pen-side interaction | 26 | 0 | 0.447 | Lactobacillus amylovorus \| PLS_N_G_C9_L24353_8281f63b | abundance |
| Host-PDS-contig abundance | post-event main effect | 26 | 0 | 0.078 | Faecousia sp034099845 \| PLS_N_G_DALL_L11624_4617ba92 | abundance |
